# A modular platform for engineering function of natural and synthetic biomolecular condensates

**DOI:** 10.1101/2021.02.03.429226

**Authors:** Keren Lasker, Steven Boeynaems, Vinson Lam, Emma Stainton, Maarten Jacquemyn, Dirk Daelemans, Elizabeth Villa, Alex S. Holehouse, Aaron D. Gitler, Lucy Shapiro

## Abstract

Phase separation is emerging as a universal principle for how cells use dynamic subcompartmentalization to organize biochemical reactions in time and space^1,2^. Yet, whether the emergent physical properties of these biomolecular condensates are important for their biological function remains unclear. The intrinsically disordered protein PopZ forms membraneless condensates at the poles of the bacterium *Caulobacter crescentus* and selectively sequesters kinase-signaling cascades to regulate asymmetric cell division^3–5^. By dissecting the molecular grammar underlying PopZ phase separation, we find that unlike many eukaryotic examples, where unstructured regions drive condensation^6,7^, a structured domain of PopZ drives condensation, while conserved repulsive features of the disordered region modulate material properties. By generating rationally designed PopZ mutants, we find that the exact material properties of PopZ condensates directly determine cellular fitness, providing direct evidence for the physiological importance of the emergent properties of biomolecular condensates. Our work codifies a clear set of design principles illuminating how sequence variation in a disordered domain alters the function of a widely conserved bacterial condensate. We used these insights to repurpose PopZ as a modular platform for generating synthetic condensates of tunable function in human cells.

## Introduction

Biomolecular condensation is a powerful mechanism underlying cellular organization and regulation in cell physiology and disease^1,2,8^. Many of these condensates are formed via reversible phase separation^2,9^, which allows for rapid sensing and response to a range of cellular challenges^10,11^. Biomolecular condensates can adopt a broad spectrum of material properties, from highly dynamic liquids to semi-fluid gels and glasses and solid amyloid aggregates^9,12–14^. Perturbing protein condensation can alter fitness^15–18^, and mutations promoting protein aggregation and other pathological phase transitions have been implicated in human disease^14,19–23^. These observations suggest that the exact material properties of a biomolecular condensate may be important for its function. However, mechanistic links between emergent properties of condensates and cellular/organismal fitness remain largely unexplored.

The bacterium *Caulobacter crescentus* reproduces by asymmetric division^24^, an event orchestrated by the intrinsically disordered Polar Organizing Protein Z, PopZ^3,4^. PopZ self-assembles into 200 nm microdomains localized to the cell poles (Fig. 1a) and forms a homogeneous membraneless compartment that excludes large protein complexes, such as ribosomes^25,26^ (Fig. 1b). In previous work, we found that retention of client proteins in the microdomain is selective for cytosolic proteins that directly or indirectly bind to PopZ, allowing for the spatial regulation of kinase-signaling cascades that drive asymmetric cell division^5^. PopZ mutants unable to condense into a polar microdomain result in severe cell division defects^27^. This well-defined and important physiological function of the PopZ microdomain makes it an ideal system to interrogate material property-function relationships *in vivo.*

**Figure 1.**
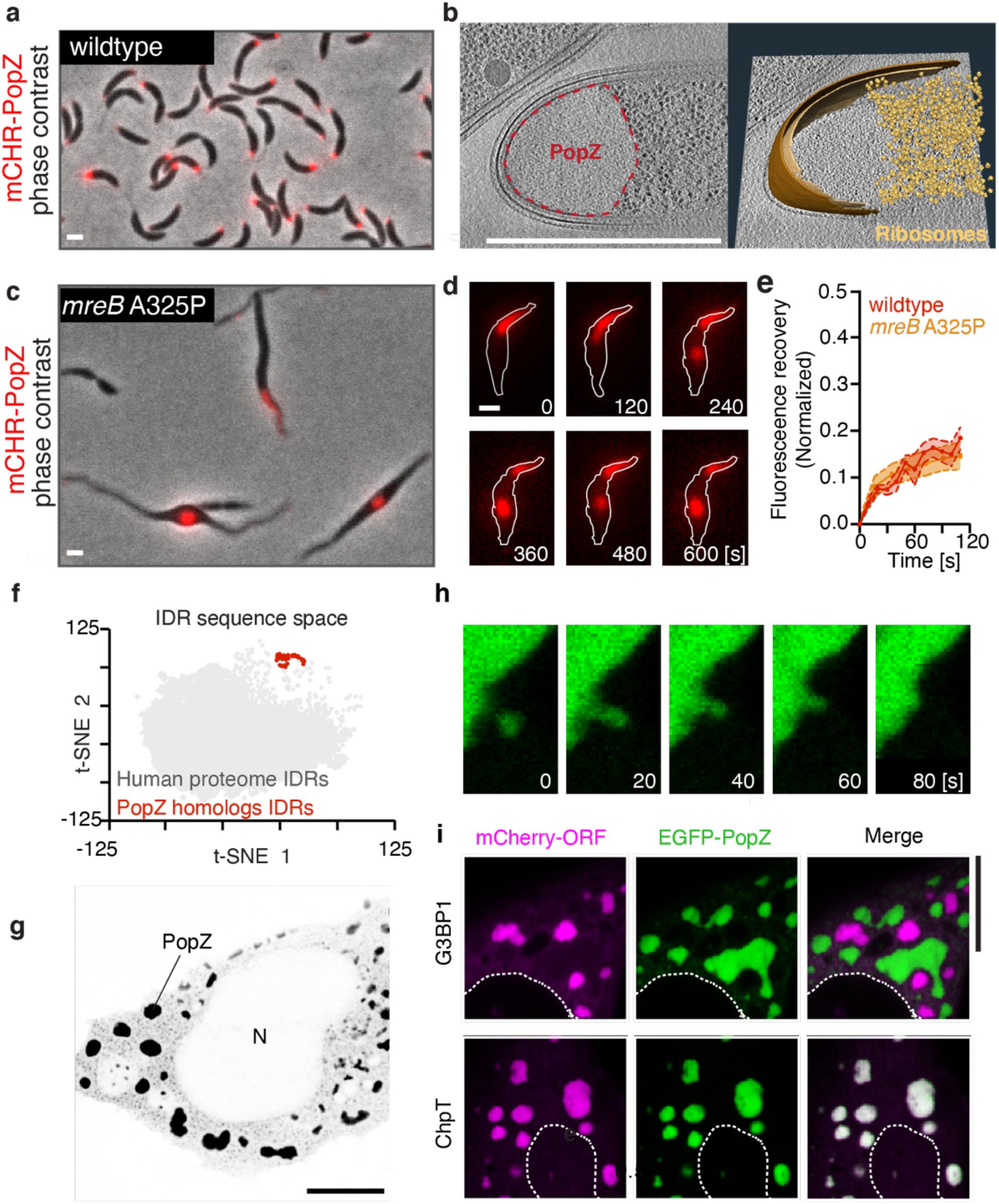
PopZ phase separates in *Caulobacter crescentus* and human U2OS cells. **a.** PopZ self-assembles at the poles of wildtype *Caulobacter* cells. A fluorescent image of ΔpopZ *Caulobacter* cells expressing mCherry-PopZ (red) from the *xylX* promoter on a high copy plasmid overlaid on a corresponding phase-contrast image. Scale bar, 1 μm **b.** The PopZ microdomain excludes ribosomes and forms a sharp convex boundary. (left) Slice through a tomogram of a focused ion beam-thinned ΔpopZ *Caulobacter* cell overexpressing mCherry-PopZ. A dashed red line shows the boundaries of the PopZ region. (right) Segmentation of the tomogram in (left) showing the outer membrane (dark brown), inner membrane (light brown), and ribosomes (gold). Scale bar, 1 μm. **c-d.** PopZ creates droplets in deformed *Caulobacter* cells. **c.** A fluorescent image of *Caulobacter* cells bearing a *mreB* A325P mutant, expressing mCherry-PopZ (red) from the *xylX* promoter on a high copy plasmid overlaid on a corresponding phasecontrast image. Scale bar, 1 μm. **d.** Fluorescent images showing the PopZ microdomain (red) extending into the cell body, concurrent with the thinning of the polar region, producing a droplet that dynamically moves throughout the cell. Frames are two minutes apart. Scale bar, 1 μm. **e.** PopZ dynamics are not affected by a release from the cell pole. Recovery following targeted photobleaching of a portion of an extended PopZ microdomain in wildtype and *mreB* A325P mutant cells. Cells expressing mCherry-PopZ from a high copy plasmid were imaged for 12 frames of laser scanning confocal microscopy following targeted photobleaching with high-intensity 561 nm laser light. Shown is the mean ± SEM of the normalized fraction of recovered signal in the bleached region; *n* equals 15 cells. **f.** IDRs of PopZ homologs cluster separately from IDRs within the human proteome. t-SNE mapping of IDR sequence composition. Each data point corresponds to the sequence composition of a single IDR. In gray are IDRs from the human proteome, and in red are IDRs from PopZ homologs within the *Caulobacterales* order. **g.** *Caulobacter* PopZ expressed in human U2OS cells forms phase-separated condensates (black) in the cytoplasm, but not the nucleus (N). **h.** *In vivo* fusion and growth of PopZ condensates in human U2OS cells. 80 seconds time-lapse images of a small PopZ condensate (green) merging with a large PopZ condensate. Scale bar, 10 μm. **i.** PopZ expressed in human U20S cells retains selectivity. (Top) EGFP-PopZ (green) and stress granule protein mCherry-G3BP1 (purple) form separate condensates. (Bottom) EGFP-PopZ (green) recruits the *Caulobacter* phosphotransfer protein mCherry-ChpT (magenta) when co-expressed in human U2OS cells. Scale bar, 10 μm.

### PopZ phase separates in *Caulobacter crescentus* and human cells

To probe the dynamic behavior of PopZ, we expressed RFP-tagged PopZ in a strain of *Caulobacter* bearing the *mreB*^A325P^ mutation^28^, which leads to irregular cellular elongation with thin polar regions and wide cell bodies^29^. While PopZ normally resides at the cell pole, in this background, the microdomain deforms and extends into the cell body before undergoing spontaneous fission, producing spherical droplets that move throughout the cell (Fig. 1c-d, Supplementary Fig. 1a). The deformation of the microdomain at the thinning cell pole and the minimization of surface tension when unrestrained by the plasma membrane provides *in vivo* evidence that the PopZ microdomain behaves as a liquid-like condensate. This observation is in line with the partial fluorescence recovery of PopZ upon photobleaching (FRAP), indicating slow internal dynamic rearrangements^5^ (Fig. 1e).

PopZ homologs are restricted to a-proteobacteria, and the sequence composition of the PopZ intrinsically disordered region (IDR) is divergent from the human disordered proteome (Fig. 1f, Supplementary Fig. 1b-e). We reasoned that human cells could serve as an orthogonal system for studying PopZ condensation. When expressed in a human osteosarcoma U2OS cell line, PopZ condensed into micron-sized cytoplasmic condensates (Fig. 1g) that underwent spontaneous fusion events (Fig. 1h) and experienced dynamic internal rearrangements, as assayed by FRAP. Importantly, even though they were expressed in human cells, PopZ condensates retained specificity for their bacterial client proteins, such as ChpT^5^, and were distinct from human stress granules (Fig. 1i). Thus, PopZ is sufficient for condensation and client recruitment, and human cells serve as an independent platform to study its behavior.

### PopZ IDR tunes the microdomain viscosity

PopZ is composed of three functional regions^27,30^ (Fig. 2a, Supplementary Fig. 2a): (i) a short N-terminal helical region (H1) used for client binding^30,31^, (ii) a 78 amino-acid (aa) IDR (IDR-78)^31^, and (iii) a helical C-terminal region (H2, H3, and H4) which is required for PopZ self-oligomerization^27^. To define the molecular mechanism driving phase separation of PopZ, we examined the contribution of each of these domains to condensation in human and *Caulobacter* cells. PopZ mutants missing either the N-terminal region (Δ1-23) or the IDR (Δ24-101) were able to form condensates in both cell types (Fig. 2b). Deletion of the IDR resulted in the formation of irregular gel-like condensates characterized by arrested fusion events in human cells (Fig. 2b) while producing dense microdomains in *Caulobacter* (Fig. 2b). In contrast, deleting any of the three predicted C-terminal helical regions (Δ102-132, Δ133-156, and Δ157-177) markedly reduced visible PopZ condensates (Fig. 2b). Therefore, the C-terminal helices are required to form condensates, and the IDR appears to play a role in tuning their material properties.

**Figure 2.**
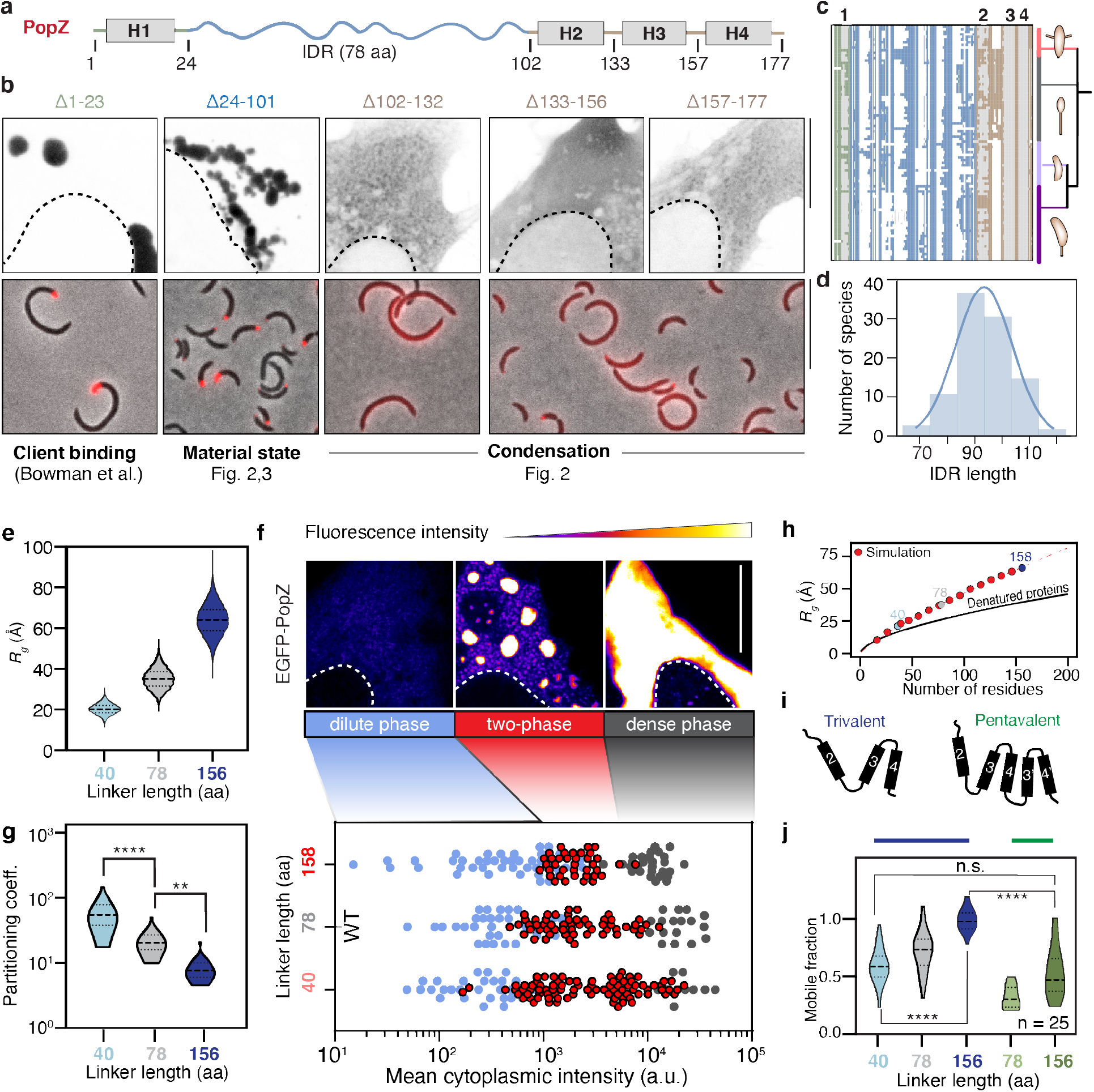
Modular organization regulates the dynamics of the PopZ condensate. **a.** Domain organization of the PopZ protein from *Caulobacter crescentus.* PopZ is composed of a short N-term region with a predicted helix, H1 (gray box), a 78 amino-acid intrinsically disordered region (IDR, blue curly line), and a C-term region with three predicted helices, H2, H3, H4 (gray boxes). **b.** Region deletion and its effect on PopZ condensation. (top) EGFP fused to five PopZ deletions (black) expressed in human U2OS cells. (bottom) mCherry fused to four PopZ deletions (Δ1-23, Δ24-101, Δ102-132, and Δ133-177) (red) expressed in *ΔpopZ Caulobacter* cells. **c.** conservation of the PopZ protein regions. Graphical representation of a multiple alignment of 99 PopZ homologs within the *Caulobacterales* order. Each row corresponds to a PopZ homolog and each column to an alignment position. All homologs encode an N-terminal region (green), an IDR (blue), and a C-terminal helical region (brown). White regions indicate alignment gaps, and gray regions indicate predicted helices 1 to 4. Phylogeny tree of the corresponding species is shown, highlighting the four major genera in the *Caulobacterales* order: *Asticcacaulis* (pink), *Brevundimonas* (gray), *Phenylobacterium* (light purple), and *Caulobacter* (dark purple). Notably, all species within the *Brevundimonas* genus code for insertion between helix 2 and helix 3. **d.** Conserved linker length within the *Caulobacterales* order. A histogram of the length of the linker of 99 PopZ homologs. The mean length is 93.6 aa with s.e.m of 1.1. **e.** Linker length and its effect on the radius of gyration. The predicted radius of gyration for a half linker (IDR-40, 40 aa) (light blue), full wildtype linker (IDR-78, 78 aa) (gray), and a double linker (IDR-156, 156 aa) (dark blue). **f.** Phase diagram of PopZ expressed in Human U2OS cells. (top) Three states of PopZ condensation: diffuse PopZ (dilute phase, blue, left), PopZ condensates (two-phase, *i.e.,* a diffused phase and condensed phase, red, middle), and a single condensate that fills most of the cytoplasm (dense phase, gray, right). A color gradient indicates EGFP fluorescence intensity from blue (low) to white (high). The nucleus boundary is shown as a white dotted line. (bottom) Phase diagrams of EGFP fused to PopZ with IDR-40, IDR-78, and IDR-156. Each dot represents data from a single cell, positioned on the x-axis as a function of the cell mean cytoplasmic intensity. The color of the dot indicates its phase, a dilute phase (blue), two-phase (red), or dense phase (gray). **g.** Quantification of the partition coefficient, *i.e.,* the ratio of the total concentration in the condensed phase to that in the protein-dilute phase, of each of the three linkers. A higher partitioning coefficient indicates denser condensates. Four (two) asterisks indicate four (two) fold difference. **h.** PopZ linker expands beyond the denatured limit. The expected R_G_ of denatured proteins as a function of the number of amino acids is shown in black^38^. Dimensions of PopZ-like linkers with varying lengths are shown in red, and dimensions of IDR-40, 78, and 156 are shown in shades of blue and gray. The red dashed line is an analytical fit with a scaling value of 0.80 with a prefactor value of 1.14. **i.** Schematics of the oligomerization domain of the wildtype PopZ (trivalent, left) and an oligomerization domain with increased valency consisting of five helices, with a repeat of helices 3 and 4 (pentavalent, right). **j.** Balance between condensation promoting and counteracting phase separation tunes condensate material properties. FRAP, shown as mobile fractions, the plateau of the FRAP curves, for PopZ with its wildtype oligomerization domain (trivalent) and a linker of three different lengths (blue and gray), as well as PopZ with an extended oligomerization domain (pentavalent) with IDR-78 (light green) and IDR-156 (dark green).

Intriguingly, the architecture of the PopZ protein from *Caulobacter crescentus* is conserved not only within the *Caulobacterales* order (Fig. 2c), but also across all a-proteobacteria (Supplementary Fig. 2b). Further, despite showing little sequence conservation, the IDR length exhibits a narrow distribution in *Caulobacterales* with a mean of 93 ± 1 aa (Fig. 2d), while other clades of a-proteobacteria occupy different length distributions (Supplementary Fig. 2c). To characterize the organization and function of the PopZ linker, we performed all-atom simulations. We found the linker adopts an extended conformation, with a radius of gyration (R_G_) of 34.4 ± 4.8 Å and an apparent scaling exponent (ν^app^) of 0.7 (Fig. 2e, Supplementary Fig. 3a). Consistent with previous studies^32–35^, the strong negative charge leads to a self-repulsing polyelectrolyte, driving expansion beyond the denatured limit and to a tight coupling between the linker length and its global dimensions (Fig. 2h). These results suggest that IDR length is constrained across species.

To test the effect of altering IDR length on its condensation, we generated PopZ mutants with a truncated or expanded IDR; namely, IDR-40, corresponding to half the wildtype IDR length and an IDR-156, corresponding to double the length of the wildtype IDR. We then tested their ability to form condensates in human cells. First, we mapped an EGFP-PopZ phase diagram as a function of concentration and IDR length.

For any liquid-liquid phase separating protein, condensates emerge when the protein concentration exceeds the saturation concentration *(C_sat_).* At a total protein concentration below *C_sat_,* the protein is uniformly dispersed (dilute phase), while above *C_sat_* demixing leads to the formation of coexisting dense and a dilute phase (two-phase regime). As the total protein concentration increases and exceeds a second threshold *(C_D_)*, the system can shift completely to the dense phase regime characterized by a single large droplet that occupies the intracellular space^9,36,37^. We found that in human cells, PopZ can exist in any of these three regimes as a function of its cytoplasmic concentration (Fig. 2f). Halving the PopZ IDR length (IDR-40) decreased *C_sat_* and increased the *C_D_*, compared to wildtype PopZ. In contrast, doubling the PopZ IDR length (IDR-156) increased *C_sat_* and decreased *C_D_*, resulting in a narrower two-phase window (Fig. 2f). Finally, increasing the IDR length decreased PopZ partitioning (Fig. 2g) and increased FRAP dynamics (Fig. 2j) in human cells. Collectively, our data suggest that the material properties of PopZ condensates are dependent on the length of its IDR.

Given the IDR offers one means of tuning PopZ material properties, we asked if altering the degree of multivalency could be used as an orthogonal control parameter. We increased the valency of the C-terminal region containing three helices (trivalent) by repeating the last highly conserved helix-turn-helix motif (Fig. 2c), resulting in PopZ variants carrying five C-terminal helices (pentavalent) (Fig. 2i). We found that pentavalent PopZ condensates had strongly reduced FRAP dynamics compared to wildtype trivalent PopZ. Combining the long IDR-156 with a pentavalent oligomerization domain (OD) normalized the FRAP dynamics to a physiological range (Fig. 2j). Taken together, our work reveals a modular design with two independent functional regions (IDR, OD) through which we can tune the material properties of the PopZ condensate, providing robust design principles for synthetic engineering of customizable condensates.

### The net charge and the charge distribution of the IDR are conserved and tune the material properties of the PopZ condensate

In addition to conserved length (Fig. 2d), the PopZ IDR shows conservation of its strong enrichment for acidic and proline residues across *Caulobacterales,* with –0.28 net charge per residue and prolines constituting 29% of the IDR residues (Fig. 3a). To test whether amino acid content plays a role in the viscosity of the PopZ microdomain, we substituted acidic residues for asparagine and proline residues for glycine. Decreasing the negative charge of the linker reduced condensate fluidity in human cells while substituting prolines for glycines slightly increased condensate fluidity (Fig. 3b). The tight conservation of some amino acid properties and the dependence of condensate material state on those properties imply that material state may be important for function.

**Figure 3.**
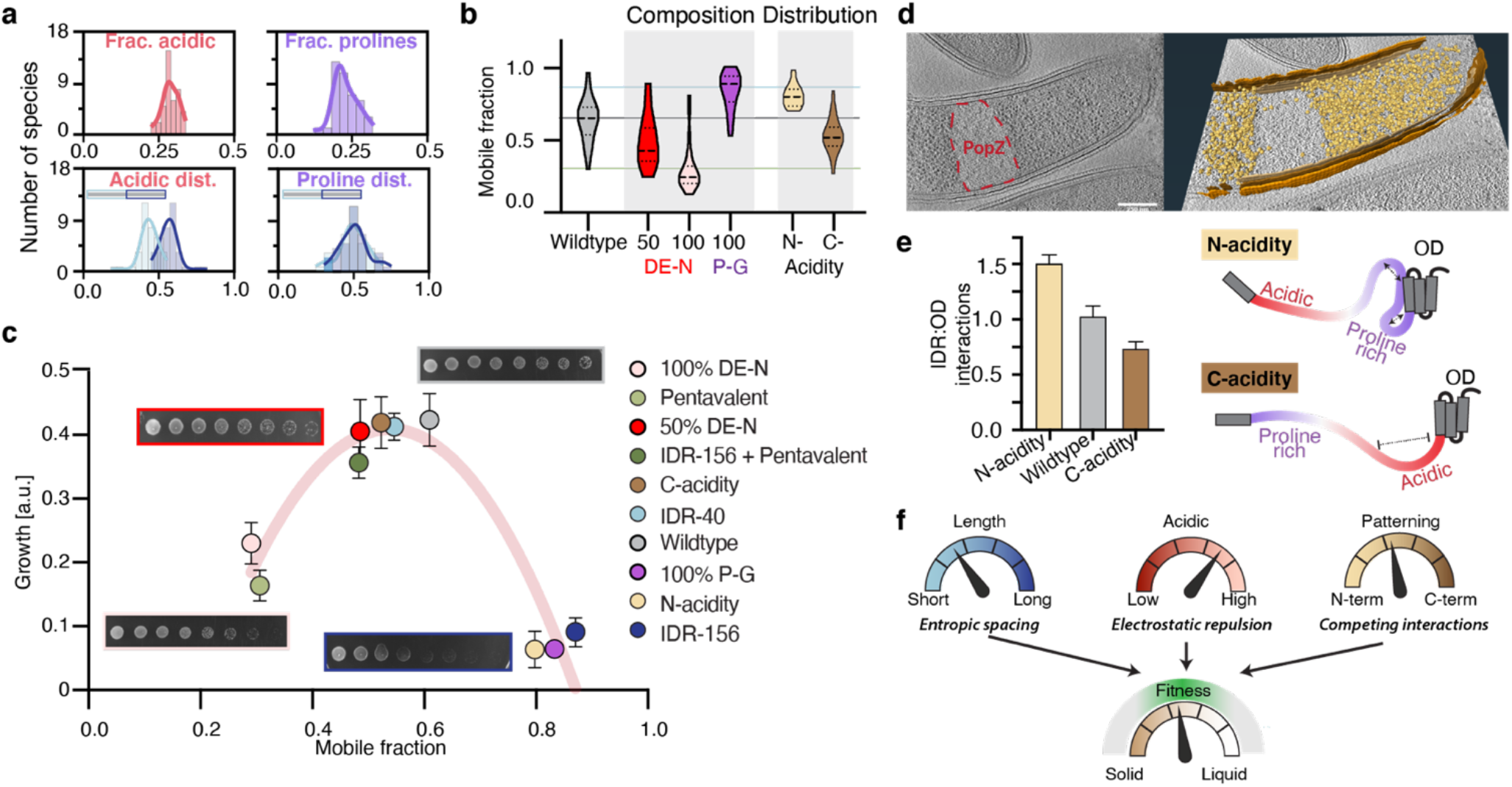
PopZ material properties are directly linked to *Caulobacter* viability and are modulated by conserved IDR properties. **a.** The sequence composition of the PopZ IDR is conserved across *Caulobacterales.* Histograms are calculated across 99 PopZ homologs within the *Caulobacterales* order and show a tight distribution for the following four parameters. (top, left) The mean fraction of acidic residues is 0.29 ±0.004 (red). (top, right) The mean fraction of prolines is 0.23±0.006 (purple). (bottom, left) Among the acidic residues within the IDR, the fraction of those found in the N-terminal half (blue, 0.57±0.011) and the C-terminal half of the IDR (orange, 0.43±0.011). (bottom, right) Among the IDR prolines, the fraction of those found in the N-terminal half (blue, 0.5±0.015) and the C-terminal half of the IDR (orange, 0.5±0.015). **b.** Amino acid composition plays a role in PopZ fluidity. FRAP, shown as mobile fractions, for PopZ with its wildtype IDR (light gray) and five mutants: Substituting either half or all of the acidic residues for asparagine (DEN 50% in red and DE-N 100% in pink, respectably), substituting all prolines for glycines (P-G 100% in purple), and moving all acidic residues to either the N-terminal part or the C-terminal part of the linker (yellow and brown, respectably). **c.** Growth is linked to PopZ material state. Growth, derived from serial dilution growth assay (Methods), as a function of FRAP mobility for ten mutants (labels indicated on the right). Examples of serial dilutions are shown for wildtype (gray box), 50% DE-N (red box), IDR-156 (dark blue box), and 100% DE-N (pink box). A polynomial fit with an R-square of 0.86 is shown in red. **d.** PopZ IDR-156 condensates retain ribosome exclusion. (left) Slice through a tomogram of a cryo-focused ion beam-thinned Δ*popZ Caulobacter* cell overexpressing mCherry-PopZ with IDR-156. (right) Segmentation of the tomogram in (left) showing annotated S-layer (orange), outer membrane (dark brown), inner membrane (light brown), and ribosomes (gold). Scale bar, 0.25 μm. **e.** Competition between intra and inter PopZ interactions. (left) Plotted is the percentage of PopZ conformations with IDR/OD interactions over an all-atoms simulation trajectory of either the wildtype PopZ (gray), PopZ with acidic residues at the N-terminal end of the linker (yellow), or PopZ with acidic residues at the C-terminal end of the linker (brown). (right) schematics of both PopZ mutants and a model suggesting that transient intramolecular IDR-OD interactions interfere with intermolecular interactions between ODs in trans, weakening the cohesive interactions that ultimately determine the material properties of the condensate. **f.** Evolutionary conserved knobs that tune PopZ material properties are linked to fitness. Chart illustrating the three identified IDR sequence features (IDR length in blue, IDR composition in red and acidity partitioning in yellow) and their modulation of PopZ material properties (solid to liquid, brown). These material properties, in turn, are linked to cell fitness (green).

We next expressed the different PopZ IDR mutants (linker length, proline, and acidity content) in Δ*popZ Caulobacter* cells and found that FRAP dynamics were consistent between *Caulobacter* and human cells (Supplementary Fig. 3b). Assaying *Caulobacter* growth in wildtype and these mutant backgrounds revealed that optimal fitness was achieved for the wildtype PopZ, while moving to either too solid or too fluid properties reduced fitness (Fig. 3c). This is particularly notable given the trivalent OD design, linker length, proline content, and acidity content are conserved (Fig. 2b, Fig. 3a). Importantly, while the pentavalent design led to more solid-like condensates in cells that reduced fitness, combining a pentavalent OD with the fluidizing double linker led to additive compensatory changes that give rise to wildtype like material properties and fitness (Fig. 3c).

Examining the behavior of the condensates, we found that unlike wildtype cells, cells expressing solid condensates did not form a second condensate at the other pole during cell growth and division, which is crucial for establishing asymmetry (Supplementary Fig. 3a,c,d). In contrast, in cells expressing more liquid condensates, like the long IDR-156, polarity was lost. The condensates were not maintained at the poles and diffused throughout the cell (Supplementary Fig. 3e). Correlative cryo-electron tomography data revealed that IDR-156 condensates retain their ability to form a barrier that excludes ribosomes (Fig. 3d, Supplementary Fig. 4). Thus, IDR-156 dynamics led to a constant reorganization of the cytosol and aberrant cell division. Collectively, our data reveal that too solid-like or too fluid-like microdomains are non-functional, suggesting that the function of the PopZ microdomain is tightly linked to its material properties, which have been precisely tuned to meet the cell’s needs. Moreover, given the importance of exclusive bipolar localization of PopZ microdomains to the progression of the *Caulobacter* cell cycle, we speculate that cells’ inability to properly localize too solid and too fluid condensates underlies-in part-their non-functionality. As the valency of the OD can restore IDR length phenotypes and *vice versa,* we suggest that a tight balance of opposing forces mediated by the IDR and the OD define this physiological window.

Since drastically changing the amino acid composition may affect several linker properties at once, we evaluated the role of potentially conserved primary sequence features. This allows us to explicitly test an alternative hypothesis - that the highly-charged IDR functions as a solubility tag, penalizing phase separation as a function of length. Accordingly, we constructed 17 scrambled versions of the IDR and measured their FRAP dynamics in human cells (Supplementary Fig. 5a,b). We calculated primary sequence features for all of these mutants (Supplementary Fig. 5c, Methods) and performed regression analysis to test which combination of features best explains the measured FRAP dynamics (Supplementary Fig. 4d). We found that a combination of differential N-versus C-acidity and differential proline enrichment best predicted experimental data with an R-square of 0.86 (Supplementary Fig. 5c,d). Notably, the values of the features used in the regression model show a narrow distribution across *Caulobacterales*, despite large differences in the actual primary IDR sequence (Supplementary Fig. 5e).

We examined two scrambled sequences that mirror one another. The C-acidity scramble has all of its acidic residues segregated to the C-terminal end of the linker, while the N-acidity scramble is simply the reverse sequence with all of the acidic residues segregated to the N-terminal end of the linker. Changing the position of the acidic residues gave rise to less dynamic or more fluid PopZ condensate compared to the wildtype protein (Fig. 3b). Strikingly, similar to our observations for IDR length and composition mutants, the FRAP dynamics of these scrambled IDR mutants correlated with cellular fitness (Fig. 3c). Because PopZ condensation is driven by OD-OD interactions (Fig. 2a), we wondered if the relative position of the acidic residues in the linker could modulate OD-OD interactions. We performed all-atom simulations on wildtype PopZ, PopZ with N-acidity, and C-acidity linkers and calculated the degree of interactions between the IDR and the OD. We found that the N-acidity IDR tends to interact more with its adjacent OD, compared to the wildtype IDR, while the C-acidity IDR tends to interact less with its adjacent OD (Fig. 3e). These findings suggest that competing IDR-OD and OD-OD interactions can regulate the dynamics of PopZ condensates.

Our results show that the function of the PopZ microdomain is tuned by its material properties, which in turn are dictated by the IDR composition and organization (Fig. 3f). By dissecting the molecular grammar of the PopZ IDR and the OD, we propose that the PopZ material properties can be explained by a molecular push-pull strategy. The valency of the OD drives condensation, while the electrostatic repulsion of the IDR fluidizes the condensates. Moreover, we show that three IDR features can be tuned to alter its repulsive nature. While IDR length and charge drive linker extension, local variations in IDR acidity can promote competing IDR-OD interactions. By subsequently testing an array of carefully designed mutants, we provide, to our knowledge, the first direct evidence that condensate material properties can gradually tune organismal fitness. Looking at the evolutionary landscape of PopZ, we find evidence suggesting that tunable IDR properties may be under selective pressure, and therefore could have helped the boom in phenotypic and ecological diversity among α-proteobacteria.

### An engineered PopTag phase separates into cytoplasmic condensates with tunable material properties

The simple modular domain architecture of PopZ, with an N-terminal client binding domain, and discrete domains that tune and drive phase separation, highlights a novel topology that is distinct from most of the currently characterized phase separating proteins (Fig. 4a). Because PopZ condensates did not interfere with human membraneless organelles such as stress granules (Fig. 1i) and seemed well-tolerated by human cells, we harnessed PopZ to engineer this simple design into a modular platform for the generation of designer condensates. We isolated the oligomerization domain and found it to be sufficient to drive condensate formation in human cells (Supplementary Fig. 6a). This “PopTag” is a C-terminal protein tag of only 76 amino acids. Just as was the case for PopZ, the material properties of these condensates could be tuned by the addition of a spacer (Supplementary Fig. 6b). To functionalize these designer condensates, we fused the PopTag to different “actor” domains. For example, by fusing the PopTag to a druginducible degron, we generated condensates whose temporal expression is under tight pharmacological control (Supplementary Fig. 6c). We also encoded biochemical reactions into these designer condensates. Fusing the PopTag to well-folded enzymes led to their condensation in the cytoplasm (Fig. 4b). To assay whether such enzymes would retain activity inside these droplets, we used TurboID, an engineered biotinylating enzyme^39^. Treating cells with biotin resulted in the biotinylation of these TurboID-PopTag condensates, as assayed by streptavidin staining (Fig. 4b), demonstrating that PopTag-generated condensates facilitate the assembly of enzymatic microreactors.

**Figure 4.**
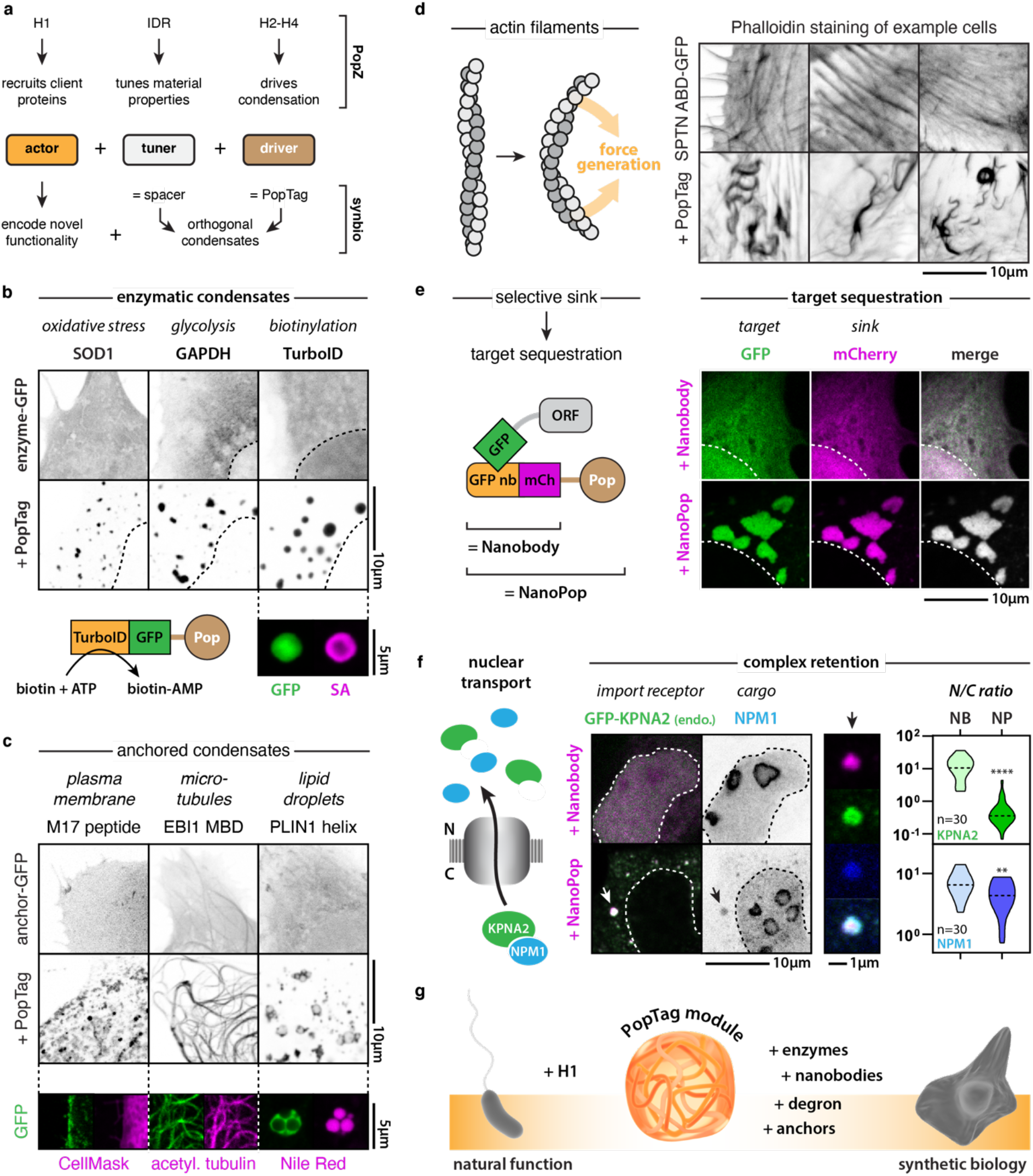
An engineered PopTag expressed in human cells phase separates into cytoplasmic condensates with tunable material properties. **a.** Re-engineering PopZ as a modular platform for the generation of designer condensates. The PopTag drives phase separation, the spacer tunes material properties, and the actor domain determines functionality. **b.** The PopTag fusion allows the condensation of enzymes. Turbo-ID maintains biotinylation activity within PopTag condensates, indicated by the biotin signal inside PopTag condensates after the addition of biotin to the cell medium. Biotin was detected by streptavidin (SA) staining. **c.** Subcellular anchors control PopTag condensate localization. Each anchor peptide is fused to EGFP in the presence or absence of the added PopTag. The M17 peptide is derived from HIV Gag protein, the microtubule-binding domain (MBD) is from EBI1, and the amphipathic helix is from PLIN1. CellMask labels the plasma membrane (purple), acetylated tubulin the microtubules (purple), and Nile Red lipid droplets (purple). Peptide-EGFP fused to PopTag is shown in green. **d.** Condensation of the PopTag on actin filaments by fusion to the spectrin actin-binding domain fusion (SPTN ABD) drives coalescence, buckling, and bending of actin filaments (phalloidin staining). **e.** NanoPop is the fusion of the PopTag to an EGFP-targeting nanobody, which allows the recruitment of EGFP (-tagged proteins) into condensates. Nanobody and NanoPop are labeled with mCherry. **f.** NanoPop condensates trap the endogenously EGFP-tagged KPNA2 in the cytoplasm of HAP1 cells, together with its client cargo protein NPM1. Violin plots show the quantification of nucleo-cytoplasmic (*N/C*) ratios. *n* is the number of cells. NB = Nanobody, NP = NanoPop. Mann-Whitney; ** p-value = 0.01, **** p-value = 0.0001. **g.** Scheme highlighting how different actor domains drive PopZ/PopTag function in nature or synthetic biology.

Accumulating data indicate that cellular condensates are spatially regulated and can interact with other subcellular structures and compartments^1,40^. To test whether our designer condensates would be amenable to such specific subcellular localization, we fused the PopTag to different “cellular anchors” - tethering the condensates in the plasma membrane, on microtubules, or on the surface of lipid droplets (Fig. 4c). Moreover, when we target the PopTag to the actin cytoskeleton by fusing it to the beta spectrin-derived actin-binding domain, the straight actin bundles of the cytoskeleton deformed and buckled, while this was not observed when we expressed the actin-binding domain by itself (Fig. 4d). These results suggest that cytoplasmic condensates can exert force upon the cytoskeleton, akin to nuclear bodies interacting with the genome^41^ and TIS-granules embedded between endoplasmic reticulum tubules^42^. These different chimeric fusions highlight the versatility of the PopTag, which can facilitate the construction of designer condensates that can differentially localize, compartmentalize biochemical reactions, or exert forces on cellular structural elements.

PopZ uses its N-terminal region to specifically recruit client proteins^30^. We replaced the N-terminal helix (H1) with an EGFP-targeting nanobody (Fig. 4e) to create “NanoPop”. These NanoPop condensates were able to efficiently sequester EGFP or EGFP-tagged proteins into cytoplasmic condensates (Fig. 4e, Supplementary Fig. 6d). As a proof-of-concept study to test whether such designer condensates can recapitulate specific cellular processes, we focused on the role of protein phase separation in nucleocytoplasmic transport. Nuclear import is mediated by importins, a class of karyopherins that bind to and facilitate the transition of cargo proteins through the nuclear pore complex (Fig. 4f). Karyopherins have also been found to bind biomolecular condensates *in vitro^43,44^.* It was recently shown that the formation of stress granules coincides with nuclear import defects, presumably due to the disruption of karyopherin availability^45,46^. While cellular stress is normally a transient event, persistent nuclear import dysregulation has been implicated in several neurodegenerative disorders^47,48^. A key pair of unanswered questions is whether the cytoplasmic retention of karyopherins is sufficient to impede nuclear transport and if retention is a direct consequence of karyopherin interaction with cytoplasmic condensates or an indirect effect of cellular stress. To answer these questions, we used NanoPop condensates to test whether the binding of karyopherins to synthetic cytoplasmic condensates is sufficient to block nuclear import of the client protein NPM1. Therefore, we endogenously tagged the karyopherin KPNA2 with EGFP in a human Hap1 cell line. Expressing EGFP-NanoPop in these cells resulted in the recruitment of KPNA2 to cytoplasmic condensates and its subsequent nuclear depletion, with a concomitant decrease in nuclear NPM1 import. In contrast, when the nanobody was expressed alone, no such defects were observed (Fig. 4e). Beyond simply reducing cargo import, NPM1 was recruited to the NanoPop condensates, showing that we were able to sequester intact complexes of client-transporter. Thus, our synthetic condensates are sufficient to drive nucleocytoplasmic transport defects in a karyopherin-dependent manner. This proof-of-concept experiment shows that tunable and functionalizable designer condensates provide a new means to untangle the contributions of specific molecular events to biological and pathological processes.

### Conclusions

As IDRs code for only 4% of bacterial proteomes, unlike 30-50% of eukaryotic proteomes^49^, their role in bacteria physiology has been largely overlooked. With accumulating evidence for the abundance of biomolecular condensates in bacterial cells^50^, and the vital role IDRs play in their formation^51^, the importance of these proteins is gaining appreciation. Bacterial IDRs differ from their eukaryotic counterparts, not only in proteome abundance but also in amino acid composition^49,52^ (Supplementary Figure 1c). These differences open new possibilities to characterize bacterial IDRs and ultimately use them to engineer synthetic biomolecular condensates to better control phase behavior in eukaryotic cells.

Here we studied the biophysical properties of the intrinsically disordered protein PopZ from the bacterium *Caulobacter crescentus*. We previously showed that PopZ forms membraneless condensates at the poles and selectively sequesters kinase-signaling cascades to regulate asymmetric cell division^3–5^. We found that PopZ self-condenses by liquid-liquid phase separation *in vivo* both in *Caulobacter* and human cells (Fig. 1). We further showed that unlike most other phase-separated IDPs, the disordered region of PopZ is not used to drive phase separation; instead, it modulates the material properties of the condensate. We found that a short structured helical domain is necessary and sufficient for phase separation (Fig. 2). We defined specific IDR characteristics that can be used to alter material properties. These include the IDR length, fraction of prolines and acidic residues, as well as the distribution of the acidic residues in the sequences (Fig. 3). Finally, we identified the optimal emergent property, a viscous liquid PopZ condensate, that is required to maintain fitness (Fig. 2,3).

Our studies have revealed a simple modular biomolecular platform, consisting of client recognition, tuner, and driver modules, enabling the engineering of a virtually unlimited set of designer condensates for synthetic biology (Fig. 4g). Given the widespread role of biomolecular condensates in cell biology, such a platform will significantly aid our understanding and manipulation of the underlying organizational principles, ultimately allowing disease modeling and the development of novel therapeutic approaches.

## Materials and Methods

### Caulobacter experiments

#### Plasmids and strains construction

Plasmids, strains, and primers are listed in Supplementary Table 2.

##### Plasmids

AP211 (pBXMCS-2 mCherry-PopZ) was amplified with primer pair 1 to remove the PopZ IDR. IDR-40 was synthesized as a gBlock gene fragment (IDT) and inserted into the linearized AP211 by Gibson assembly^53^ to make pKL539. pKL540 and pKL577 were constructed in a similar fashion with primer pairs 2 and 3 and gBlock gene fragments that codes for IDR-156 and H3-H4, respectively. To make pKL581, pKL540 was amplified with primer pair 3, and gBlock H3-H4 was inserted into the linearized pKL540 by Gibson assembly. To make pKL699-704, AP211 was digested with KpnI and SacI. Corresponding gBlocks were inserted into the digested and linearized AP211 by Gibson assembly. PCRs were performed with the KOD Hot-start 2× master mix (Novagen), and cloning was performed using Gibson Assembly 2× Master Mix (New England BioLabs, NEB) following the manufacturer’s instructions. The sequence of each insert was verified by Sanger sequencing (Sequetech).

##### Strains

To make KL6212, purified plasmid pBXMCS-2 mCherry-PopZ from AP211 cells was electroporated into KL5943. To make all other strains, purified plasmids were transformed into Δ*popZ* cells by electroporation and plated on marked PYE plates. The resulting colonies were screened for mCherry fluorescence after induction with xylose and confirmed by western blots.

### Diffraction-limited data collection and image analysis

Images were collected using a Leica DMi8 S microscope equipped with a Hamamatsu C9100 EM-CCD camera, a 100x oil-immersion objective (1.63 NA), and a SPECTRA X light engine (Lumencor). Cell outlines and intensity profiles were identified using MicrobeJ^54^ and manually filtered to eliminate false positives. Custom MATLAB (The MathWorks) scripts were used to calculate the average fluorescence intensity profile along the long axis of the cell.

### Fluorescence recovery after photobleaching

Photobleaching experiments were performed using an LSM710 line-scanning confocal microscope (Zeiss) with a 60x oil immersion objective with a numerical aperture (NA) of 1.4. A circular region of interest (ROI) within a PopZ microdomain was bleached using a high-intensity 561 nm laser and 50% bleaching power. Pictures were taken at the rate of five per minute for three minutes. Control pictures (cells and background) were taken under the same conditions. Normalization and photobleaching corrections were performed^55^.

### Serial dilution plating viability assay

Strains were grown in M2G with appropriate antibiotics to an OD600 of 0.3. Ten microliters of each dilution were spotted onto PYE plates in triplicates. Plates were incubated at 30°C for two days and imaged with Gel Doc XR Imaging System (BioRad). The mean density for each spot was calculated using FIJI following background subtraction. The growth value for each strain was defined as the mean density at the sixth dilution divided by the mean density at the first dilution. The parabolic fit was conducted using Prism.

### Bioinformatics

PopZ homologs were identified based on the C-term region using BLAST^56^, and taxonomy was extracted from NCBI. A phylogenetic tree was determined based on full-length sequences using Geneious Prime 2020.0.4 (https://www.geneious.com). NetSurfP-2.0^57^ was used to detect intrinsically disordered regions, and JPred^58^ to detect secondary structures in the full-length homologs. Custom python scripts were used for regression analysis and visualization.

### All-atom simulations

All-atom simulations were run using the ABSINTH implicit solvent model and the CAMPARI Monte Carlo simulation (http://campari.sourceforge.net/)^59^. The combination of ABSINTH and CAMPARI has been used previously to effectively sample the conformational behavior of disordered proteins with good agreement to experiment, notably in the context of highly charged and highly proline-rich IDRs^32,60^. All simulations were started from randomly generated nonoverlapping random-coil conformations, with each replica using a unique starting structure. Monte Carlo simulations evolve the system via a series of moves that perturb backbone and sidechain dihedral angles along with the rigid-body coordinates of both polypeptides and explicit ions. Simulation analysis was performed using CAMPARITraj (www.ctraj.com) and MDTraj^61^. The protein secondary structure was assessed using the DSSP algorithm^62^.

ABSINTH simulations were performed with the ion parameters derived by Mao *et al.,* with the notable exception of the double linker for which an enhanced Na^+^ ionic radius (2.32 Å vs. 1.16 Å) was applied to prevent non-physiological chelation^63^. All simulations were run at 10 mM NaCl and 310 K. An overview of the simulation input details is provided in Supplementary Table 3, while a summary of simulation results is provided in Supplementary Table 4.

A major challenge in the sampling of disordered proteins reflects an effective exploration of conformational space. The highly repulsive and expanded nature of the linker provides some advantages here, in that conformational space is substantially reduced by the polyelectrolytic nature of the chain. Simulations reveal no substantial secondary structure (Supplementary Fig. 7a), with good agreement between analogous sub-regions examined in different length constructs. Further, histograms of Rg revealed a smooth distribution consistent with a well-sampled ensemble without substantial local kinetic traps (Supplementary Fig. 7c).

To assess sampling for full-length PopZ, we compared simulation-derived secondary structure profiles for wildtype PopZ, N-acidity, and C-acidity mutants (Supplementary Fig. 7b). In agreement with good conformational sampling, we observed nearly perfectly overlapping helicity profiles for the N and terminal regions that remain unchanged between the three constructs, giving us confidence that simulations are relatively converged with respect to the relevant order parameters of interest. As with the linker constructs, smooth distributions for the *Rg* are again consistent with a well-sampled conformational ensemble (Supplementary Fig. 1d).

### Human cells experiments

#### Human PopZ plasmids

PopZ and derived mutant constructs for expression in human cells were generated through custom synthesis and subcloning into the pcDNA3.1+N-eGFP backbone by Genscript (Piscataway, USA). The mCherry-G3BP1 plasmid was a kind gift of Dr. Kedersha and Dr. Anderson (Brigham and Women’s Hospital).

#### Human cell culture and microscopy

U2OS cells (ATCC, HTB-96) and EGFP-KPNA2 HAP1 cells were grown at 37 °C in a humidified atmosphere with 5 % CO2 for 24 h in Dulbecco’s Modified Eagle’s Medium (DMEM), high glucose, GlutaMAX + 10 % Fetal Bovine Serum (FBS) and pen/strep (Thermo Fisher Scientific). Cells were transiently transfected using Lipofectamine 3000 (Thermo Fisher Scientific) according to manufacturer’s instructions. Biotinylation experiments using TurboID-PopTag condensates were performed as described in detail^64^. Cells grown on coverslips were fixed for 24 h after transfection in 4 % formaldehyde in PBS. Slides were mounted using ProLong Gold antifade reagent (Life Technologies). Confocal images were obtained using a Zeiss LSM 710 confocal microscope. Images were processed using FIJI.

#### Endogenous GFP-tagging of KPNA2

To stably create cells expressing a KPNA2-EGFP fusion from their endogenous genetic locus, cells were CRISPR edited. HAP1 cells were transfected using TurboFectin 8.0 (Origene) according to the manufacturer’s instructions with a plasmid encoding spCas9 and an sgRNA targeting the C-terminal sequence of KPNA2 (5’-AGGCTACACTTTCCAAGTTC-3’). In addition, a PCR product was transfected. The product contains the sequence to restore the C-terminus of KPNA2 with a silent mutation upstream of the endogenous C-terminal PAM sequence (to prevent recutting of the edited sequence), 3xFLAG and EGFP followed by a P2A-coupled puromycin resistance gene. Following transfection, cells were incubated for 2 days and then selected over a period of 1 week with 1 μg/mL puromycin. Resistant cells were harvested and plated at a density of 0.5 cells/well in 96-well plates in 20% FBS containing medium to obtain single cell derived colonies. Colonies were grown for 2-4 weeks and when sufficiently grown, cells were washed and then lysed in Bradley lysis buffer at 56°C (10 mM Tris-HCl (pH 7.5), 10 mM EDTA, 0.5% SDS, 10 mM NaCl and 1 μg/mL proteinase K). The genomic DNA was extracted from the lysate using ethanol-salt precipitation and centrifugation. The target site, the C-terminus of KPNA2, was amplified by in-out PCR with the following primers: fwd: 5’-GAAGAATGTGGAGGCTTAGACAAAATTG-3’, rv: 5’-CAGCCATTCTCGGGCCGATC-3’. The PCR product was sequenced with reverse primers 5’-GGAACGTCGTCTCTTGTAGC-3’ and 5’-CCGTAGGTGGCATCGCCC-3’ by Sanger Sequencing (Macrogen).

#### FRAP measurements in human cells

U2OS cells were cultured in glass-bottom dishes (Ibidi) and transfected with GFP-PopZ constructs as described above. After 24 h, GFP-PopZ condensates were bleached and fluorescence recovery after bleaching was monitored using Zen software on a Zeiss LSM 710 confocal microscope with incubation chamber at 37 °C and 5 % CO2. Data were analyzed as described previously^55^. In brief, raw data were background subtracted and normalized using Excel, and plotted using GraphPad Prism 8.4.1 software.

#### Statistical analysis

All data was analyzed using Graphpad Prism 8.4.1 and Excel. Statistical tests details are shown in figure legends.

### Cryo-Electron Tomography

#### Sample preparation

Log phase *Caulobacter crescentus* (OD600 between 0.2 and 0.5) grown in M2G media were diluted 1:10 in fresh M2G media and induced for 4-5 hours with 3% xylose at 28 °C in a shaking incubator. For plunge freezing of *Caulobacter,* induced cells were placed on ice, and concentrated to an effective OD600 of 3.0 by centrifugation. For whole-cell tomography, cells were diluted to an effective OD600 of 0.2 and plunge frozen in a similar manner.

In order to reduce formation of crystalline ice, 1 uL of 50% w/v trehalose was added to 9 ul of the cell suspension immediately before plunge-freezing. 4 uL of the cell suspension were added to the carbon side of a glow-discharged Cu 200 mesh R2/1 Quantifoil grid and manually blotted from the back to remove excess liquid, and were plunge-frozen in an ethane/propane mixture cooled to liquid nitrogen temperatures using a custom-built manual plunger (Max Planck Institute for Biochemistry). Grids were clipped into an Autogrid support ring to facilitate downstream handling. The frozen grids were kept at liquid nitrogen temperatures for all subsequent steps.

#### Cryo-Fluorescence Microscopy

Frozen grids were observed with a CorrSight inverted microscope (Thermo Fisher Scientific) using EC Plan-Neofluar 5x/0.16NA and EC Plan-Neofluar 40x/0.9NA air objectives (Carl Zeiss Microscopy), a 1344×1024 px ORCA-Flash 4.0 camera (Hamamatsu), and an Oligochrome lightsource, with excitation in four different channels (405/488/561/640 nm); red (mCherry-PopZ) and green (GFP-ribosomes) were used. Data acquisition and processing were performed using MAPS 2.1 and MAPS 3.6, respectively (Thermo Fisher Scientific). After acquiring a grid map at 5X magnification, regions of interest were imaged at 40x magnification to identify cells with PopZ domains.

#### Cryo-Focused Ion Beam (FIB) Milling

Grids with *Caulobacter* were prepared using cryo-FIB milling as previously described using an Aquilos (Thermo Fisher Scientific) dual-beam SEM equipped^65,66^. Briefly, areas covered with a monolayer of cells were targeted first for coarse milling with an ion beam current of 0.10-0.50 nA, followed by fine milling using 10-50 pA. Lamella width was typically 10-12 um. Five to eight lamellae were prepared on each grid in one session, with a target thickness of ~150 nm.

#### Cryo-Electron Tomography

*Caulobacter* lamellae were visualized on a Titan Krios (Thermo Fisher Scientific) operating at 300 kV accelerating voltage with a Gatan K2 Summit camera equipped with a Quantum energy filter. Regions of interest were determined by correlating TEM and FM images. Tilt series of *Caulobacter* were obtained using SerialEM^67,68^ using both bi-directional and dose-symmetric tilt schemes^69^ over a tilt range of +/- 60°, in increments of 2° or 3°, at a pixel size of 0.4265 nm or 0.3457 nm. Each tilt image was collected using electron counting mode and with dose-fractionation. Exposure times for each tilt were adjusted to keep an approximately constant number of counts on the sensor. The cumulative dose for each tilt series was usually between 120 to 180 e/A^2^.

#### Tomogram Reconstruction

The movies corresponding to each tilt were motion-corrected using MotionCor2 software^70^. Tilt series alignment and reconstruction was done using IMOD^71–73^. Tilt series were aligned using the patch-tracking modality and reconstructed using weighted back-projection. If needed, individual tilts with excessive motion, poor contrast, or camera errors were excluded from the final reconstruction. Non-linear anisotropic diffusion (NAD) filtering was applied to tomograms in iMOD to enhance contrast for presentation in IMOD.

#### Membrane Segmentation

Membranes were detected using TomoSegMemTV^74^. Membrane and ribosome annotations were visualized with Amira (Thermo Fisher Scientific).

#### Ribosome classification

To determine the location of ribosomes within the tomogram, we used template matching, 3D alignment, and classification. Tilt series were preprocessed using Warp v1.0.9^75^ for sub-frame motion correction and 3D-CTF estimation. Tilt-series were then aligned using IMOD^71-73^ as above and imported into Warp for final reconstruction.

Template matching of ribosomes within tomograms was performed in Warp. First, 261 manually picked particles were aligned and averaged, and this structure was subsequently used as an initial template. All extracellular particles were initially discarded based on cell boundaries defined in Dynamo. Obvious false positives (e.g., membrane segments) were manually excluded. The remaining particles were used for 3D alignment and classification in Relion v3.1 ^76^. Particles were subject to successive rounds of binary classification with a large (500 Å or 83 binned pixels) mask, the smaller class which contained particles that did not appear as ribosomes were removed from subsequent rounds. This was done until the two classes reached about equal population, both clearly representing ribosomes.

Coordinates and orientations of the remaining particles were imported into Amira for visualization^77^. The few particles residing inside the PopZ domain (<1%) were visually inspected to verify their identity. Some, such as membrane-bound ribosomes, are likely true positives, while some in the middle of the cell are likely false positives. Overall, the PopZ domains dramatically exclude ribosomes compared to their concentration in the rest of the cell.

## Supplementary Data

Supplementary Data are available.

## Acknowledgments

K.L. acknowledges support from the Gordon and Betty Moore Function (award no. GBMF 2550.03) to the Life Sciences Research Foundation and the Weizmann Institute of Science National Postdoctoral Award Program for Advancing Women in Science. S.B. acknowledges an EMBO Long Term Fellowship. Work in the A.D.G. lab is supported by NIH grants R35NS097263 and R01AG064690). L.S. is a BioHub investigator, and her work is supported in part by the National Institute of General Medical Sciences, the National Institutes of Health (R35-GM118071). E.V. is supported by an NIH Director’s New Innovator Award 1DP2GM123494-01. V.L is supported by NIH 5T32GM7240-40 NIH R35GM118290 (to Susan S. Golden). This work was performed in part at the cryo-EM facility, which was built and equipped with funds from UC San Diego, an initial gift from the Agouron Institute and an NSF MRI grant (DBI 1920374), and at the San Diego Nanotechnology Infrastructure (SDNI) of UCSD, a member of the National Nanotechnology Coordinated Infrastructure, which is supported by the National Science Foundation (Grant ECCS-1542148). We thank Sergey Suslov and Thomas Laughlin for assistance with cryo-FIB milling and ribosome classification, respectively.

