## Supplementary Figures and Tables for "A modular platform for engineering function of natural and synthetic biomolecular condensates"

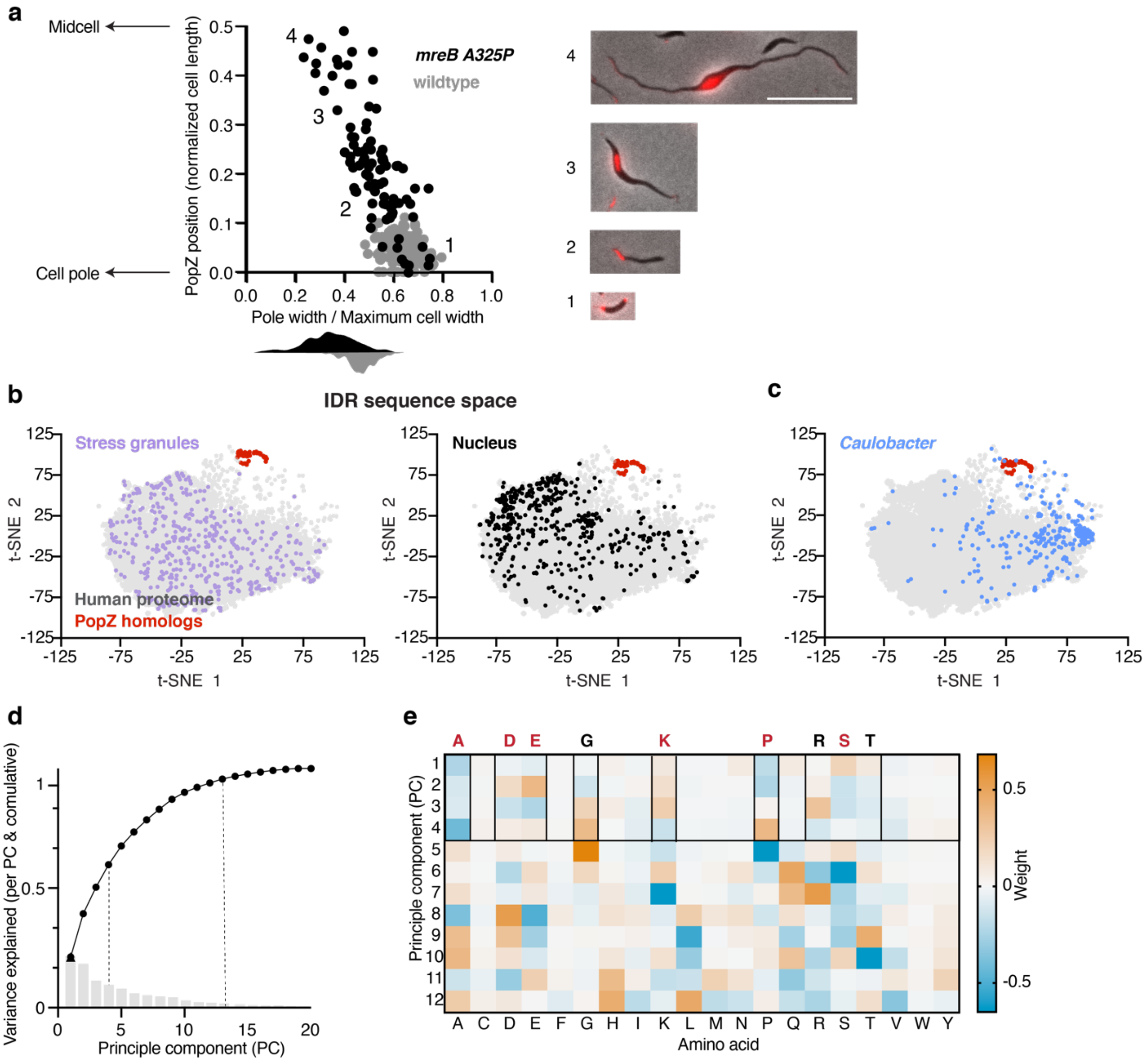


**Supplementary Figure 1. PopZ localization as a function of cell width and a comparison between IDRs in human cells and the PopZ IDR**

**a.** Localization of the PopZ microdomain as a function of the cell width. (left) Scatter plot showing the PopZ microdomain's localization as a function of the ratio between the pole width and the maximum cell width (width ratio). Each dot corresponds to a single cell. In wildtype cells, the width ratio is maintained with a mean of 0.64 and s.e.m of 0.004 (gray violin plot), while in cells expressing *mreB* A325P, the width ratio varies with a mean of 0.5 and s.e.m of 0.01. In wildtype cells, there is a slight anti-correlation (cross-correlation score of -0.16) between PopZ localization and the width ratio (gray dots). In cells expressing *mreB* A325P, PopZ appears away from the pole as the width ratio decreases, resulting from the thinning of the polar region and expansion of the cell body. This relationship is quantified by a strong anti-correlation (cross-correlation score of -0.80) between PopZ localization and the width ratio. (right) Four snapshots of cells. Scalebar 10 μm. **b.** PopZ sequence composition differs from IDRs found in human proteins that are part of either stress granules or the nucleolus. Shown are t-SNE mappings of IDR sequence composition. Each data point corresponds to the sequence composition of a single IDR. In gray are IDRs from the human proteome, and in red are IDRs from PopZ homologs within the *Caulobacterales* order. IDRs found in human stress granules proteins are shown in purple (left), and IDRs found in the nucleolus are shown in black (right). **c.** PopZ sequence composition is separate from most IDRs found in *Caulobacter crescentus*. The composition of IDRs found in Caulobacter proteins (blue) intersects with the composition of IDRs found in the human proteome (gray), unlike PopZ (red). **d-e.** Nine amino acids explain 60% of the variance in sequence composition across human and *Caulobacter* IDRs. **d.** Results of a principal component dimensionality reduction using the twenty amino acids as features. The first three (four) PCs explain 50% (60%) of the variance in the data, and 12 PCs are required to explain 95% of the variance in the dataset. **e.** Graphical representation of the 12 PCs used for the t-SNE analysis. The first four PCs use a linear combination of eight amino acids (black highlight), including polar residues (Asp, Glu, Lys, Arg), Ala, Pro, as well as Gly, Ser, and Tyr, which are enriched in IDRs of RNA binding proteins[1,2]. Amino acids that are present in PopZ IDR are highlighted in red as well.

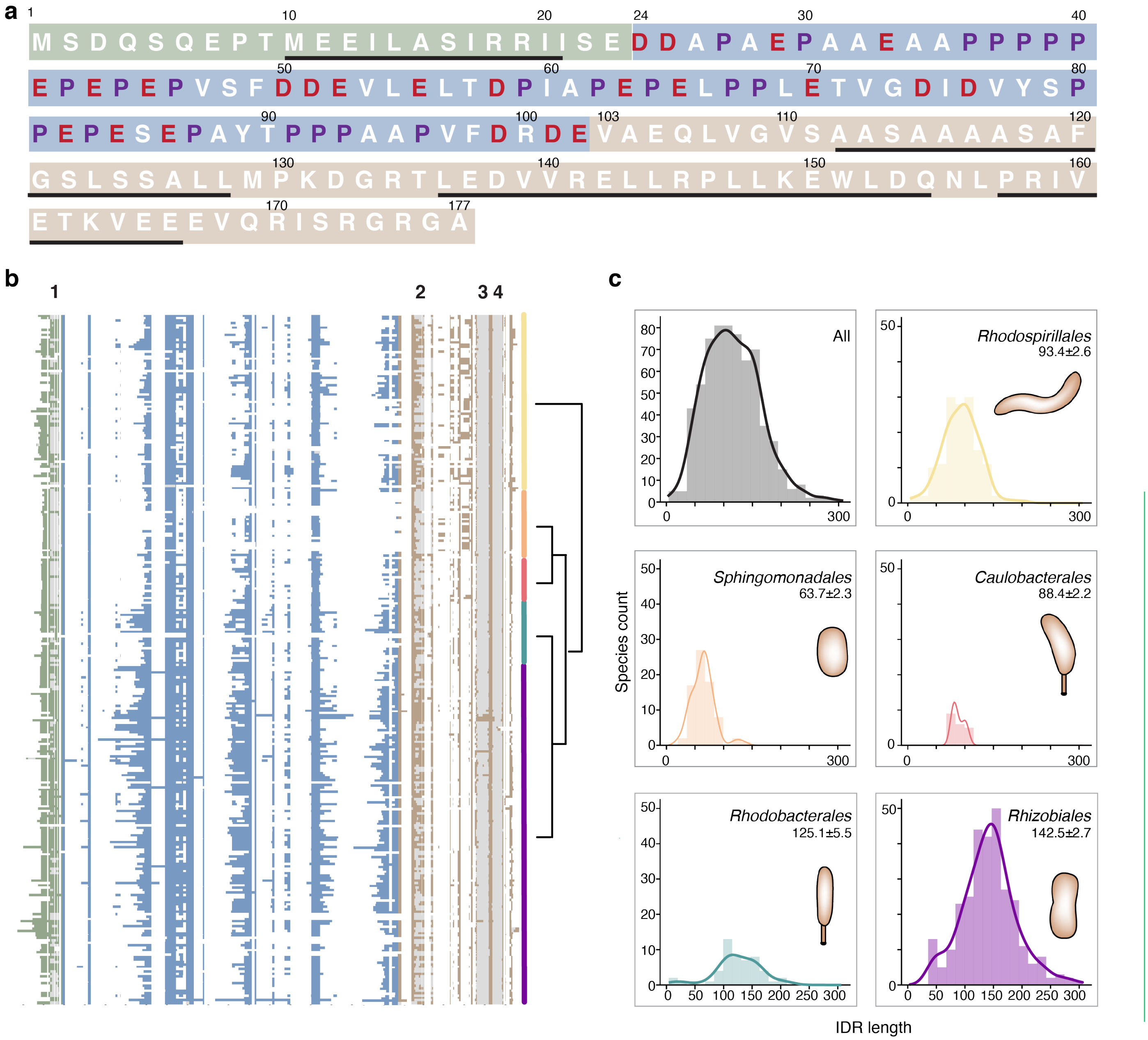


**Supplementary Figure 2. PopZ sequence across alpha-proteobacteria.**

**a.** PopZ primary sequence. The N-terminal, IDR, and C-terminal regions are indicated above using a green, blue, and brown background. Within the IDR, prolines are colored in purple and negatively charged residues in red. Black rectangles indicate the boundaries of predicted α-helices. **b.** Conservation of the PopZ protein regions within α-proteobacteria. Graphical representation of multiple alignment of 655 PopZ homologs across α-proteobacteria. Each row corresponds to a PopZ homolog and each column to an alignment position. All PopZ homologs encode a short helical N-terminal region (green), an IDR (blue), and a helical C-terminal helical region (brown). The C-terminal region is divided into two sub-modules: a region that includes helix 2, which varies in length and helicity, and a region that includes helices 3 and 4, which is highly conserved. White regions indicate alignment gaps, and gray regions indicate predicted helices 1 to 4. Phylogeny tree of the corresponding species is shown, highlighting five major orders within α-proteobacteria: *Rhodospirillales* (yellow), *Sphingomonadales* (orange), *Caulobacterales* (red), *Rhodobacterales* (green), and *Rhizobiales* (purple). **c.** Wide distribution of linker length across α-proteobacteria. Shown are the length distribution of the PopZ IDR across all of the 655 representatives α-proteobacteria and per order. Mean and s.e.m is reported for each.

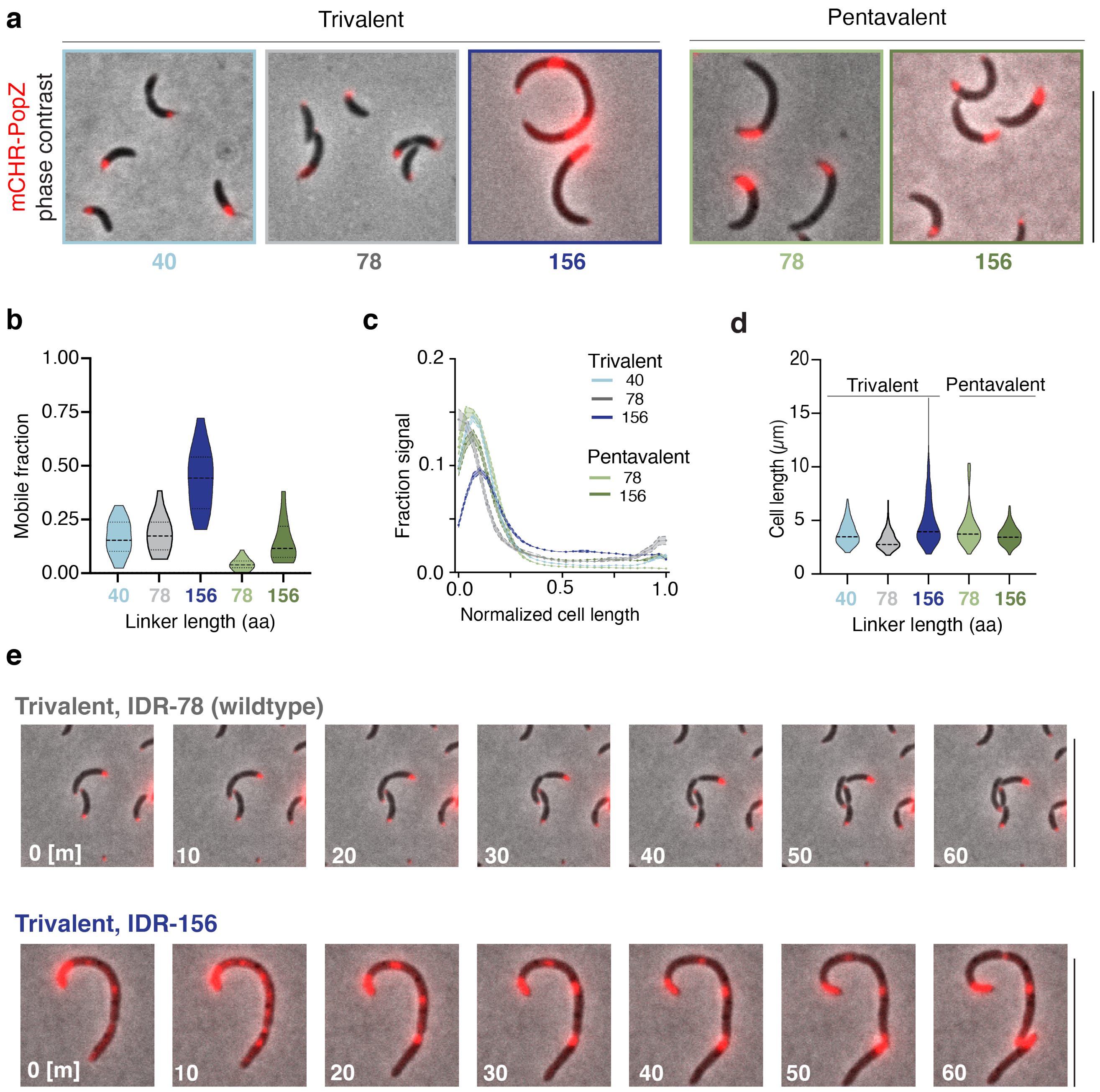


**Supplementary Figure 3. IDR length and OD valency affect PopZ localization.**

**a.** Linker length and its effect on condensate localization in *Caulobacter*. ΔpopZ *Caulobacter* cells expressing mCherry fused to PopZ with an IDR of different lengths and either a trivalent or a pentavalent c-terminal region. mCherry-PopZ with IDR-40 or the wildtype IDR-78 maintains its localization at the poles of the cell, while mCherry-PopZ with IDR-156 demonstrates condensates throughout the cytoplasm. The mutants of PopZ with pentavalent c-term both show polar localization. Scale bar, 10 µm. **b.** Balance between condensation promoting and obstructing tunes material properties. A violin plot of the distribution of FRAP measurements for the different mutants in *Caulobacter*. FRAP, shown as mobile fractions, for PopZ with its wildtype oligomerization domain (trivalent) and a linker of three different lengths (shades of blue and gray), as well as PopZ with an extended oligomerization domain (pentavalent) with IDR-78 (light green) and IDR-156 (dark green). **c.** Fluorescence intensity profiles along normalized cell length for ΔpopZ *Caulobacter* cell overexpressing different PopZ mutants (n = 51, 250, 250, 164, and 142 for trivalent OD with IDR-40, IDR-78, IDR-156, and pentavalent OD with IDR-78 and IDR-156). Cells expressing wildtype PopZ show microdomains at both poles. In cells expressing IDR-78 with pentavalent OD, a second polar PopZ microdomain is lost and recovered in IDR-156 with pentavalent OD. **d.** Cell length for the different mutants. A violin plot of the distribution of cell lengths for the different mutants. At least 30 cells were measured for each condition. **e.** Time-lapse showing 10 minutes snapshots of ΔpopZ *Caulobacter* cells expressing wildtype PopZ and IDR-156.

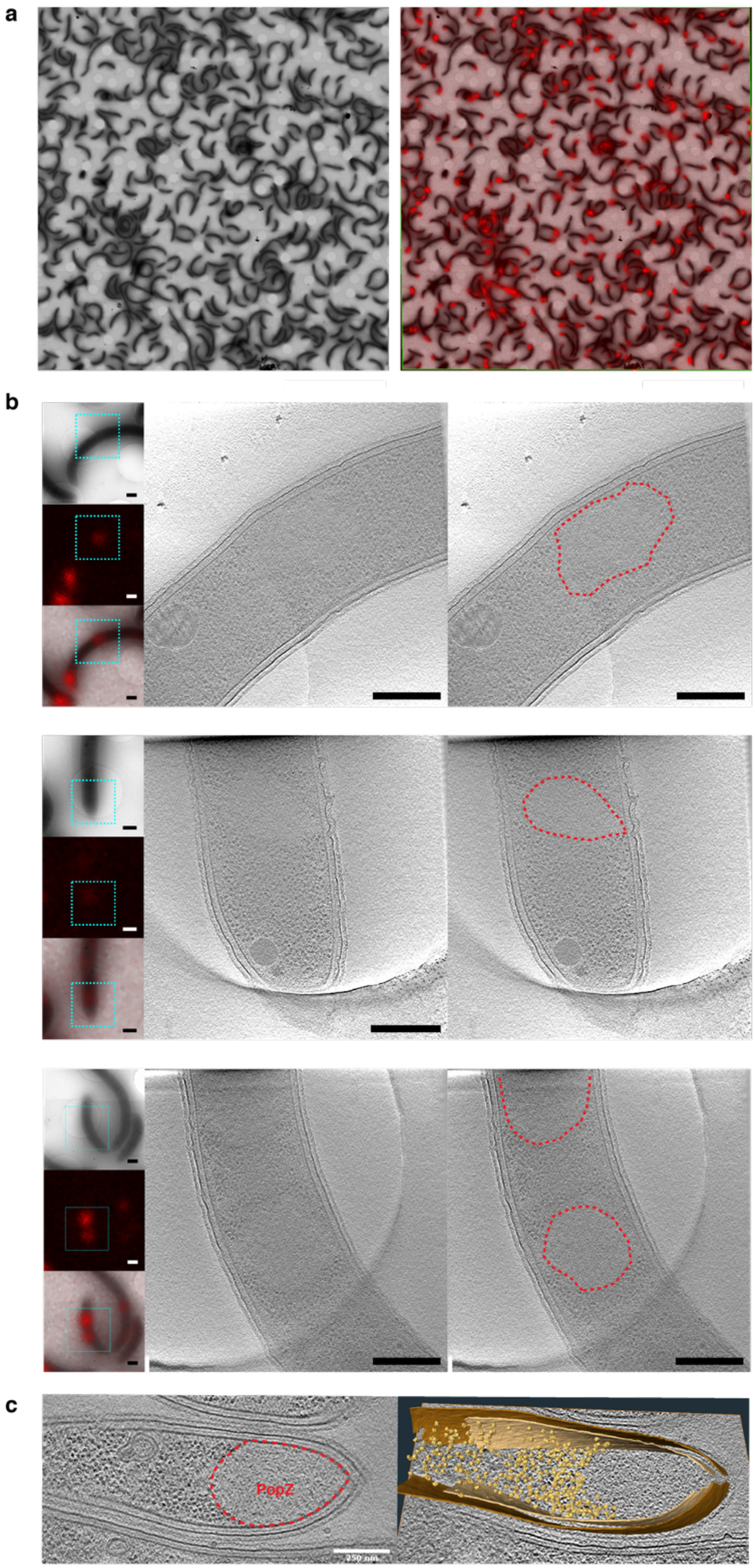

a
b
c
PopZ
250 nm

**Supplementary Figure 4. Correlative fluorescence cryo-electron tomography showing PopZ with IDR-156 excludes ribosomes**

**a-b.** Correlative fluorescence cryo-electron tomography imaging of cells expressing mCherry-PopZ with IDR-156. a. A field of view of cells on a cryo-grid visualized using an electron microscope (left) and overlaid with the corresponding cryo-fluorescence image (right). PopZ is shown in red. Scale bar, 20 µm. b. In each row shown are (left, top) a cell in low-magnification with a blue square indicating the region for high-magnification tomography imaging, (left, middle) the same low-magnification imaging area in cryo-fluorescence, and (left, bottom) an overlay of the two channels. (middle) a slice of the re-constructed tomogram showing a ribosome free region. (right) the same slice as in (middle) with the PopZ region annotated, as resolved from the fluorescence channel. Scale bar, 0.5 µm. **c.** (left) Slice through a tomogram of a cryo-focused ion beam-thinned ΔpopZ *Caulobacter* cell overexpressing mCherry-PopZ with IDR-156 and pentavalent OD. (right) Segmentation of the tomogram in (left) showing annotated outer membrane (dark brown), inner membrane (light brown), and ribosomes (gold). Scale bar, 0.25 µm.

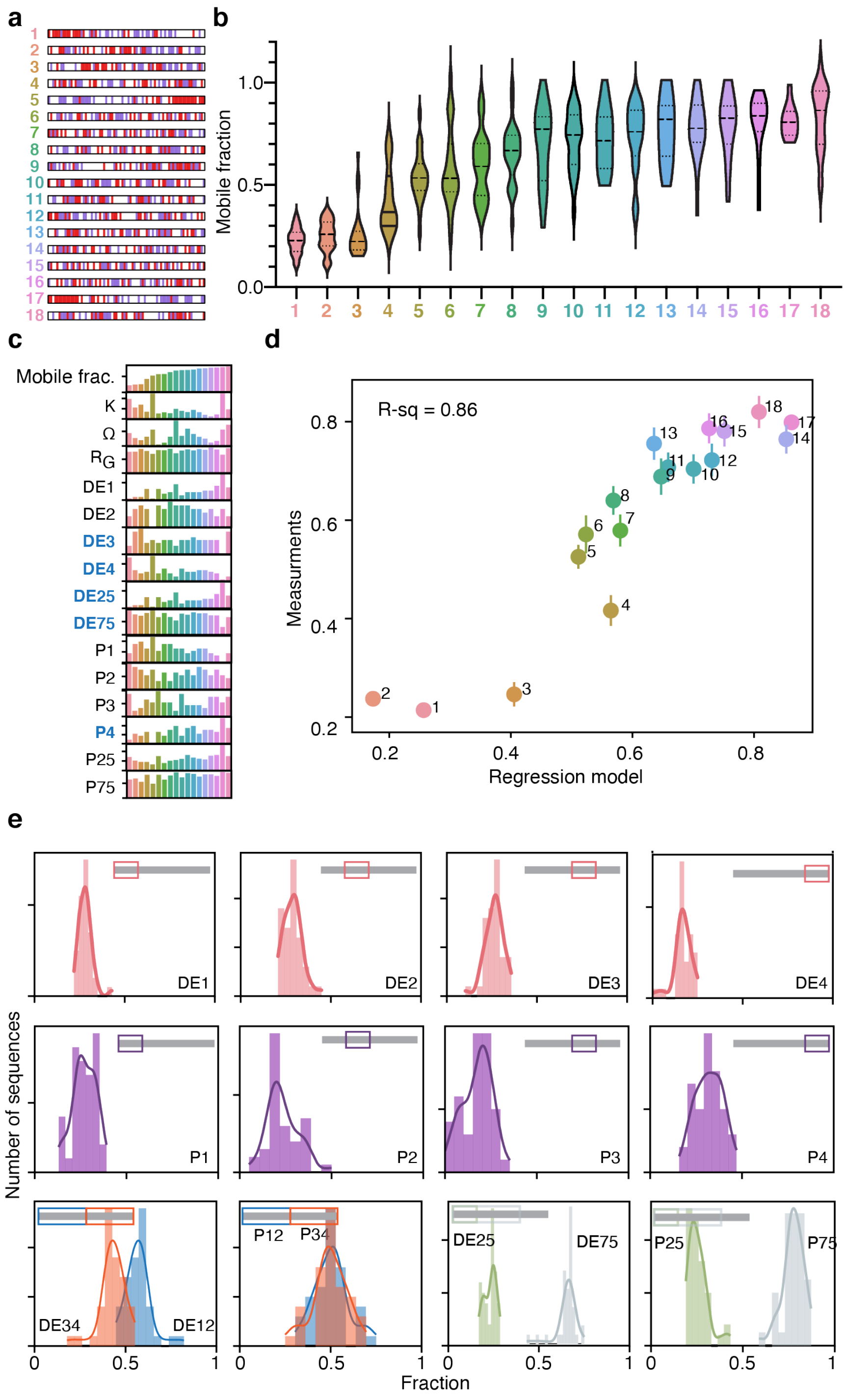


**Supplementary Figure 5. Charge distribution contributes to PopZ liquidity.**

**a.** A graphical representation of 18 scrambles of the wildtype PopZ IDR. Red indicates an acidic amino acid, and purple indicates a proline. The scrambles are sorted by their mobile fraction (b). **b.** A violin plot of FRAP measurements of PopZ condensates in human cells. The FRAP data are shown as mobile fractions, which vary from 30% (for L1), 63% (for the wildtype IDR L8), to 80% (for L18). **c.** 16 Bar plots, each is showing the values of a feature across the 18 scrambles. The bars are color-coded by the scramble numbers (a). Features include (i) FRAP mobile fraction, (ii) $0 \leq K \leq 1$ parametrizing the degree of mixing (0) *vs.* segregation (1) of oppositely charged residues[3], (iii) $0 \leq \Omega \leq 1$ parametrizing the degree of mixing (0) *vs.* segregation (1) of charged and proline residues[4], (iv) $R_g$ radius of gyration as calculated by an all-atom-simulation (Methods), (v) 0 ≤DE1-4≤ 1, the fraction of acidic residues in each quarter of the sequence, (vi) 0 ≤DE25, DE75≤ 1, N-terminal fraction that includes 25% and 75% of the IDR acidic residues, (vii) 0 ≤P1-4≤ 1, the fraction of proline residues in each quarter of the sequence, (viii) 0 ≤P25, P75≤ 1, N-terminal fraction that includes 25% and 75% of the IDR prolines. **d.** A regression model using five of 25 features derived from the IDR sequence (Methods). The model presented has an R-square of 0.86 and is a linear combination of differential N- versus C-acidity, as well as the number of prolines present in the C-terminal quarter of the IDR (DE3, DE4, DE25, DE75, P4). **e.** The features used in the regression model are conserved across *Caulobacterales*. The histograms are calculated across 99 PopZ homologs within the *Caulobacterales* order.

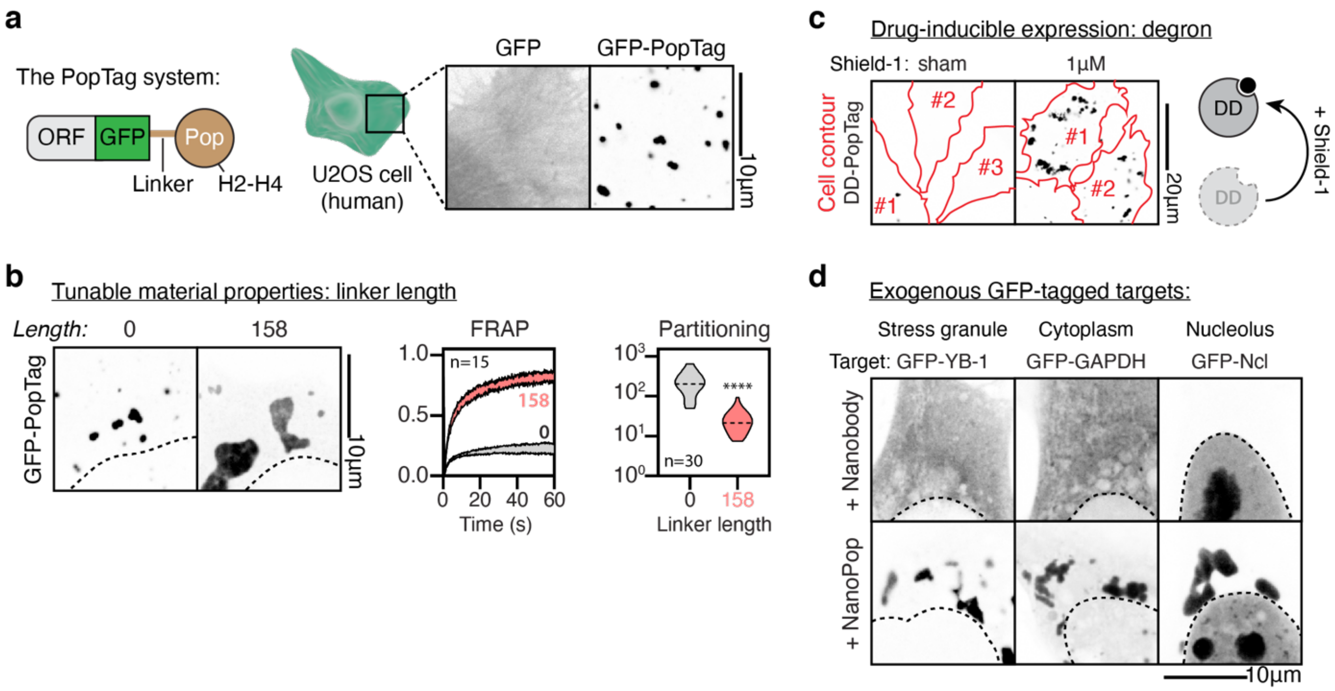


**Supplementary Figure 6. PopTag condensates have tunable functionality.**

**a.** Scheme highlighting setup of the PopTag system and formation of EGFP-PopTag condensates in U2OS cells. **b.** Changing the linker length alters the FRAP dynamics and partitioning coefficient of PopTag condensates. Student's t-test; **** p-value < 0.0001. **c.** Fusing PopTag to the drug-stabilized degron (DD) allows for the pharmacological control of PopTag expression. The addition of Shield-1 stabilizes the degron and prevents degradation of DD-PopTag condensates. **d.** NanoPop condensates can sequester different EGFP-tagged client proteins upon transient expression.

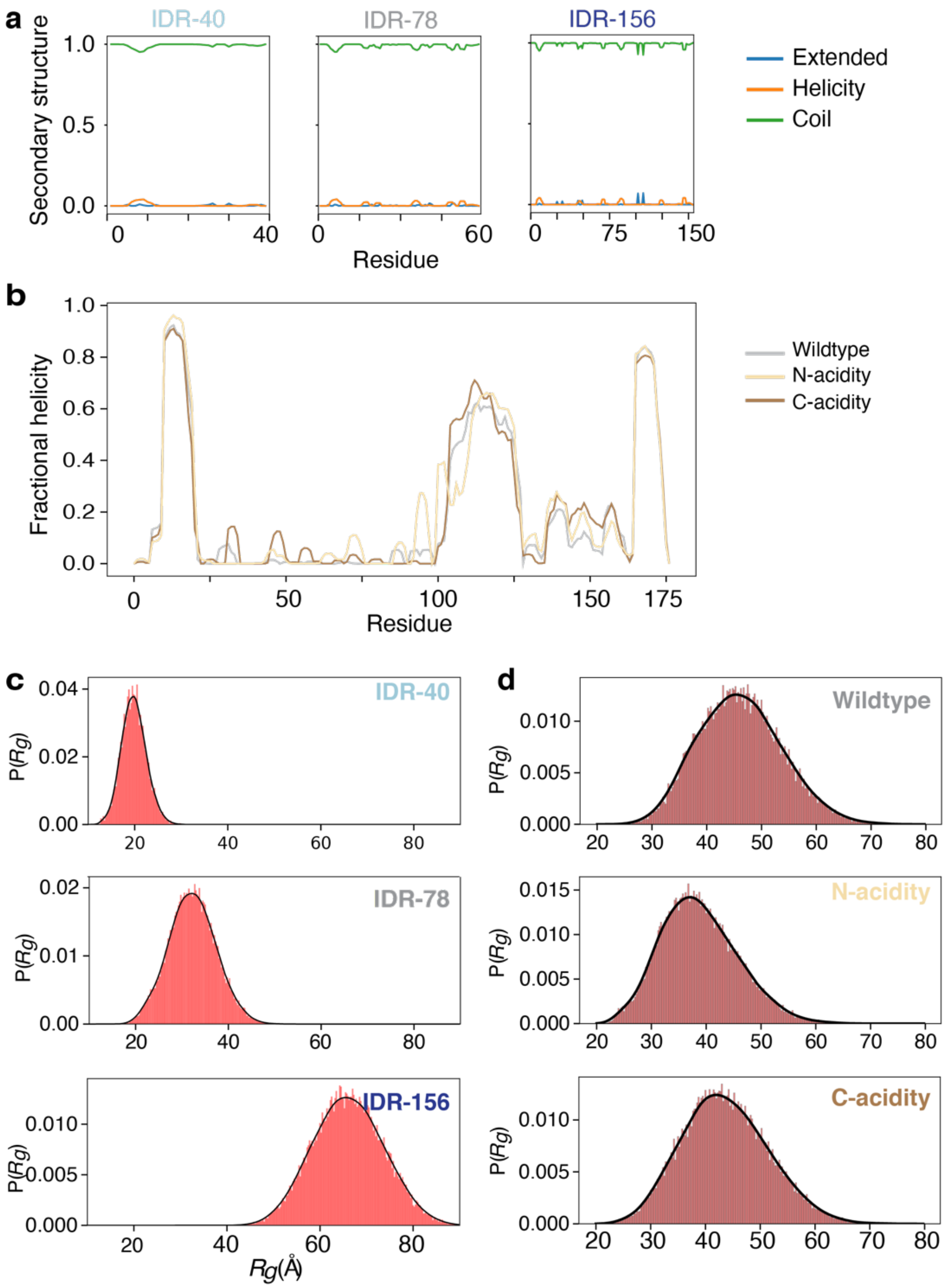


**Supplementary Figure 7. Simulations show complete sampling**

**a.** Secondary structure analysis for the IDR-40 (half), IDR-78 (full), and IDR-156 (double) PopZ linkers reveals no significant helicity or beta-sheeted (extended) structure. **b.** Comparative helicity analysis for WT PopZ and the N-acidity and C-acidity mutants. **c.** Distributions for the radii of gyration of IDR-40 (half), IDR-78 (full), and IDR-156 (double) PopZ linkers. The broad, smooth distributions match expectations for a flexible polymer suggest robust and effective conformational sampling. **d.** Histograms for the radius of gyration of WT PopZ and the two linker charge mutants.

### Supplementary Tables

#### Supplementary Table 1. Linker sequences

| Linker ID | Sequence |
|---|---|
| L1 | FIPRSDAYAAVGLAPTPPTPFPLDEPAPPLP-<br>SPPPVIDTEPYPVAPASPAEPEPVAEPDDEDAEEEEPEDDEVDE |
| L2 | ALPPPRVPLSAEEPVPPPLTPPLPAAPPPDDPIPEAEDEEPDVFDDEAEAP-<br>FSYDTGEVEIPADPAPPPEEEYSVDA |
| L3 | LPEPIFPPPYFEIPPSPPVPPEDETDPEARPPPVDPAGLDTPPDAESADDEVEEPADEEEEA-<br>SPTYVLAPPADLAPVA |
| L4 | LESVPPDEPPPEDPETFDAPPAPAEVPIDDEVPAEASGAEPPASLPPPVRPYLDE-<br>AETIYPPPEFDPADEPDLEPVA |
| L5 | PPIFAVPPPPYLAPPPDPPPPPYPASAGFPSVPVTTAPRLPVPEPSIPAE-<br>AAEADEPATLDVEEEEDEEDDEDDLEDE |
| L6 | LEPVDSVPPDSDEYPYDPEFETPVAPEDLVLEPLDAEGPFPPPPADDPVEIEPPA-<br>PAAEAPDEASPPTPIEPPAAER |
| L7 | PAPVVAPPVDPPPAPDDETPEPVDIPVPPALEAPELDFPEAYYPEGP-<br>PAADDRLPTEPEESAPDSAIFELESDPEPE |
| L8, IDR-78 | DDAPAEPAAEAAPPPPPEPEPEPVSFDDEVLELTDPIAPEPELPPLETVG-<br>DIDVYSPPEPESEPAYTPPPAAPVFDRD |
| L9 | AYEPPEPDEDPEEDPDPEGALPEPDEPPEAPDAPSLEIFPTEVDPYVALEVAAFAT-<br>LASAPRPPDVPVPEIPSPPDP |
| L10 | PAASVIYDAPSIEEFPPPPEDGEAAEPPPALPDEVTPVSPEVEDFAPPPP-<br>PAEDPPRPAYEDDEPPAVLEEPDTTLDL |
| L11 | PFVATDPEPALEEIAPTDYSDPAVPLPDPPEDDPRFEPVAAAASADVPPEEDPVPELPEP-<br>SEEEPPPDPEGILPPAY |
| L12 | RDAEAVPFYPPELVELPEPEPPPTAVDVPESDDPAVIAGATETEEELPEPID-<br>SPSPALPPAFPPPADPDPPPDEYDEA |
| L13 | PPSYPDDSEIEPAAAFDPPEPPEPLPLPDGALEPDSPPPADEAAPDIPVEVREATPVPE-<br>AAPEDEPEDPVTFPEYLV |
| L14 | APEEAPPDTEIYTPEPSPEYPRPPVPEPDPTEPVDAVSESVLPLAVPEEALPAIED-<br>AGPDALPDDAPDPEPPEFPAFD |

| | |
|---|---|
| L15 | DFPESPGDVVADTPYEVPEPAEAPDVLEPPSELYPERAPDPPAE-AAPEPPPDTADATEFPAEPVEPLDPPLSDIIEPP |
| L16 | DLDLLAEPEAPDEEPREAEAPAPEPGPTFPEPITVAPESPDVPPPPADAPLESDIPEPAPYD-YEEPVPPVDADFSVP |
| L17 | EDELDDEDDEEDEEEEVDLTAPEDAEAAEAPISPEPVPLRPATTVPVSPF-GASAPYPPPPPDPPPALYPPPPVAFIPP |
| L18 | EVVSIAPDLDPEEDPDPDALPAPSATTVLIDPEPEAGRE-PEAFEDLPPPPEDPDPEAFPYAPVYEPPEPPPEVAPSA |
| IDR-40 | DDAPAEPAAEAAPPPPPEPEPEPVSFDDEVLELTDPIAPE |
| IDR-156 | DDAPAEPAAEAAPPPPPEPEPEPVSFDDEVLELTDPIAPEPELPPLETVG-DIDVYSPPEPESEPAYTPPPAAPVFDRDDDAPAEPAAEAAPPPPPEPEPEPVSFDDEVLELTDPIAPEPELPPLETVGDIDVYSPPEPESEPAYTPPPAAPVFDRD |

**Supplementary Table 2. Caulobacter strains, plasmids, and primers**

| Plasmids | | | |
|---|---|---|---|
| **Number** | **Description** | **Strain number** | **Source** |
| AP211 | pBXMCS-2, mCherry-PopZ. High copy plasmid (kanr) | AP211 | [5] |
| pKL539 | pBXMCS-2, mCherry-PopZ with IDR-48 | KL6254 | This study |
| pKL540 | pBXMCS-2, mCherry-PopZ with IDR-156 | KL6252 | This study |
| pKL577 | pBXMCS-2, mCherry-PopZ with IDR-78 and pentavalent OD | KL6322 | This study |
| pKL581 | pBXMCS-2, mCherry-PopZ with IDR-156 and pentavalent OD | KL6326 | This study |
| pKL699 | mCherry-PopZ with L5 | KL6607 | This study |
| pKL700 | mCherry-PopZ with L17 | KL6531 | This study |
| pKL702 | pBXMCS-2, mCherry-PopZ with IDR 100% P-G | KL6529 | This study |
| pKL704 | pBXMCS-2, mCherry-PopZ with IDR 100% DE-N | KL6530 | This study |

| | | | |
|---|---|---|---|
| pKL703 | pBXMCS-2, mCherry-PopZ with IDR 50% DE-N | KL6554 | This study |

| Strains | | |
|---|---|---|
| **Number** | **Description** | **Source** |
| KL5820 | Δ*popZ* | LS5130[6] |
| KL5943 | *mreB*::A325P*mreB* | JAT702[7] |
| KL6212 | AP211 in *mreB*::A325P*mreB* | This study |
| KL6256 | PopZ with IDR-40 in Δ*popZ*. | pKL539 electroporated into KL5820 |
| KL6261 | PopZ with IDR-156 in Δ*popZ*. | pKL540 electroporated into KL5820 |
| KL6341 | mCherry-PopZ with IDR-78 and pentavalent OD in Δ*popZ*. | pKL577 electroporated into KL5820 |
| KL6349 | mCherry-PopZ with IDR-156 and pentavalent OD in Δ*popZ* | pKL581 electroporated into KL5820 |
| KL6561 | mCherry-PopZ with L5 in Δ*popZ* | pKL699 electroporated into KL5820 |
| KL6545 | mCherry-PopZ with L17 in Δ*popZ* | pKL700 electroporated into KL5820 |
| KL6541 | mCherry-PopZ with all prolines replaced with glycines in Δ*popZ* | pKL702 electroporated into KL5820 |
| KL6542 | mCherry-PopZ with all acidic residues replaced with asparagines in Δ*popZ* | pKL704 electroporated into KL5820 |
| KL6562 | mCherry-PopZ with half of all acidic residues replaced with asparagines in Δ*popZ* | pKL703 electroporated into KL5820 |

| Pair number | Forward | Reverse | Used to make plasmid |
|---|---|---|---|
| 1 | GGGACGCGGCGCCTAA-GAATTCCTGCAGCCCGGGGG | GGTTCTTGAGACTGATCG-GACATGGTAC-CATGCATATTAATTAAGGCGCCTGC | pKL539 |
| 2 | CCTGTGTTTGAC-CGTGATGCCGAG-CAGCTGGTCGGC | GGCTGGTGCATCATCCTCCGA-GATGATGCGTCGAATGGAGG | pOpto540 |
| 3 | CGTGGAGCTTAAGAATTCCTG-CAGCCCGG | CGTCCACGAGAGATACGCTGCAC | pOpto577, pOpto581 |

**Supplementary Table 3. Overview of simulation input settings**

| System | Droplet radius (Å) | Total number of replicas | Steps per simulation | Equilibration steps | Conformations per simulation | Total ensemble size |
|---|---|---|---|---|---|---|
| WT linker | 122 | 30 | 60 000 000 | 6 000 000 | 1200 | 36 000 |
| Half linker | 81 | 5 | 27 000 000 | 2 000 000 | 1250 | 6 250 |
| Double linker | 184 | 30 | 120 000 000 | 6 000 000 | 1200 | 34 500 |
| Full protein | 200 | 30 | 150 000 000 | 10 000 000 | 1500 | 45 000 |
| Full (N terminal charge block) | 200 | 30 | 150 000 000 | 10 000 000 | 1500 | 45, 000 |
| Full (C terminal charge block) | 200 | 30 | 150 000 000 | 10 000 000 | 1500 | 45 000 |
| Sequence length titrations | Variable | 20 | 66 000 000 | 4 000 000 | 1320 | 39, 600 |

**Supplementary Table 4. Summary of simulation details for all-atom simulations.**

| System | $N_{res}$ | Mean $R_g$ (Å) | Mean $R_e$ (Å) | $\nu^{app}$ |
|---|---|---|---|---|
| Wildtype linker | 78 | 32.4 | 90.6 | 0.72 |
| Half linker | 39 | 19.9 | 49.3 | 0.61 |
| Double linker | 154 | 66.2 | 182.8 | 0.78 |
| Full protein | 177 | 46.3 | 107.7 | 0.64 |
| Full (N terminal charge block) | 177 | 38.9 | 92.5 | 0.58 |
| Full (C terminal charge block) | 177 | 43.8 | 96.7 | 0.65 |

The apparent scaling exponent ($\nu^{app}$) was calculated by fitting a linear model to the intra-residue distances, as described previously[8]. The fitting of $\nu^{app}$ becomes systematically less useful and appropriate as a chain deviates from homopolymer behavior (as has been extensively discussed previously) and as such specific values are relatively uninformative for the full-protein constructs[9].
